# EZH2-TTP-mTORC1 Axis Drives Phenotypic Plasticity and Therapeutic Vulnerability in Lethal Prostate Cancer

**DOI:** 10.1101/2025.08.07.669104

**Authors:** Beatriz German, Katherine L. Morel, Teia Noel, Nadia Boufaied, Deborah L. Burkhart, Sujun Chen, Felipe Dezem, Xintao Qui, Henry W. Long, Stefan DiFazio, Sylvan Baca, Ayesha A. Shafi, Matthew L. Freedman, Himisha Beltran, Christopher J. Sweeney, Housheng Hansen He, Myles Brown, Jasmine T. Plummer, Simon R.V. Knott, David P. Labbe, Leigh Ellis

**Affiliations:** Center for Prostate Disease Research, Murtha Cancer Center Research Program, Department of Surgery, Uniformed Services University of the Health Sciences, Bethesda, MD, USA; The Henry M. Jackson Foundation for the Advancement of Military Medicine, Inc., Bethesda, MD, USA; Genitourinary Malignancies Branch, Center for Cancer Research, National Cancer Institute, Bethesda, MD, USA; South Australian Immunogenomics Cancer Institute, University of Adelaide, Adelaide SA, Australia; Cedars-Sinai Center for Bioinformatics and Functional Genomics, Los Angeles CA, USA; Cancer Research Program, Research Institute of the McGill University Health Center, Montreal, QC, Canada; Department of Cancer Biology, Dana-Farber Cancer Institute, Harvard Medical School, Boston MA, USA; West China School of Public Health, West China Fourth Hospital, and State Key Laboratory of Biotherapy, Sichuan University, Chengdu, China; Department of Developmental Neurobiology, St. Jude Children’s Research Hospital, Memphis TN, USA; Center for Functional Cancer Epigenetics, Dana-Farber Cancer Institute, Boston MA, USA; Department of Medical Oncology, Dana-Farber Cancer Institute, Harvard Medical School, Boston MA, USA; Princess Margaret Cancer Centre, University Health Network, Toronto, ON, Canada; Department of Medical Biophysics, University of Toronto, Toronto, ON, Canada; Department of Biomedical Sciences, Cedars-Sinai Medical Center, Los Angeles CA, USA; Department of Anatomy and Cell Biology, McGill University, Montreal, QC, Canada

**Keywords:** EZH2, PI3K, mTOR, Phenotypic Plasticity, Translation

## Abstract

Phenotypic plasticity is a recognized mechanism of therapeutic resistance in prostate cancer (PCa), however current knowledge of driver mechanisms and therapeutic interventions are limited. Using genetically engineered mouse models (GEMMs) devoid of Pten and Rb1, we previously demonstrated the chromatin reprogramming factor enhancer of zeste homolog 2 (EZH2) as an important regulator of alternative transcription programs promoting phenotypic plasticity. Here, using a multi-omics approach we demonstrate that EZH2 regulates multilineage cell states dependent on the RNA binding protein Tristetraprolin (TTP) that mediates RNA stability and activation of translation. Combined chemical inhibition of EZH2 and PI3K/mTORC1 resulted in superior anti-tumor activity in murine and human phenotypic plastic models and was most significant when this combination was used with castration or enzalutamide. Together, these data indicate phenotypic plasticity dependence on coordination between EZH2, TTP and mTORC1 signaling that represent novel therapeutic dependencies for this lethal PCa phenotype.

**Graphical abstract:** 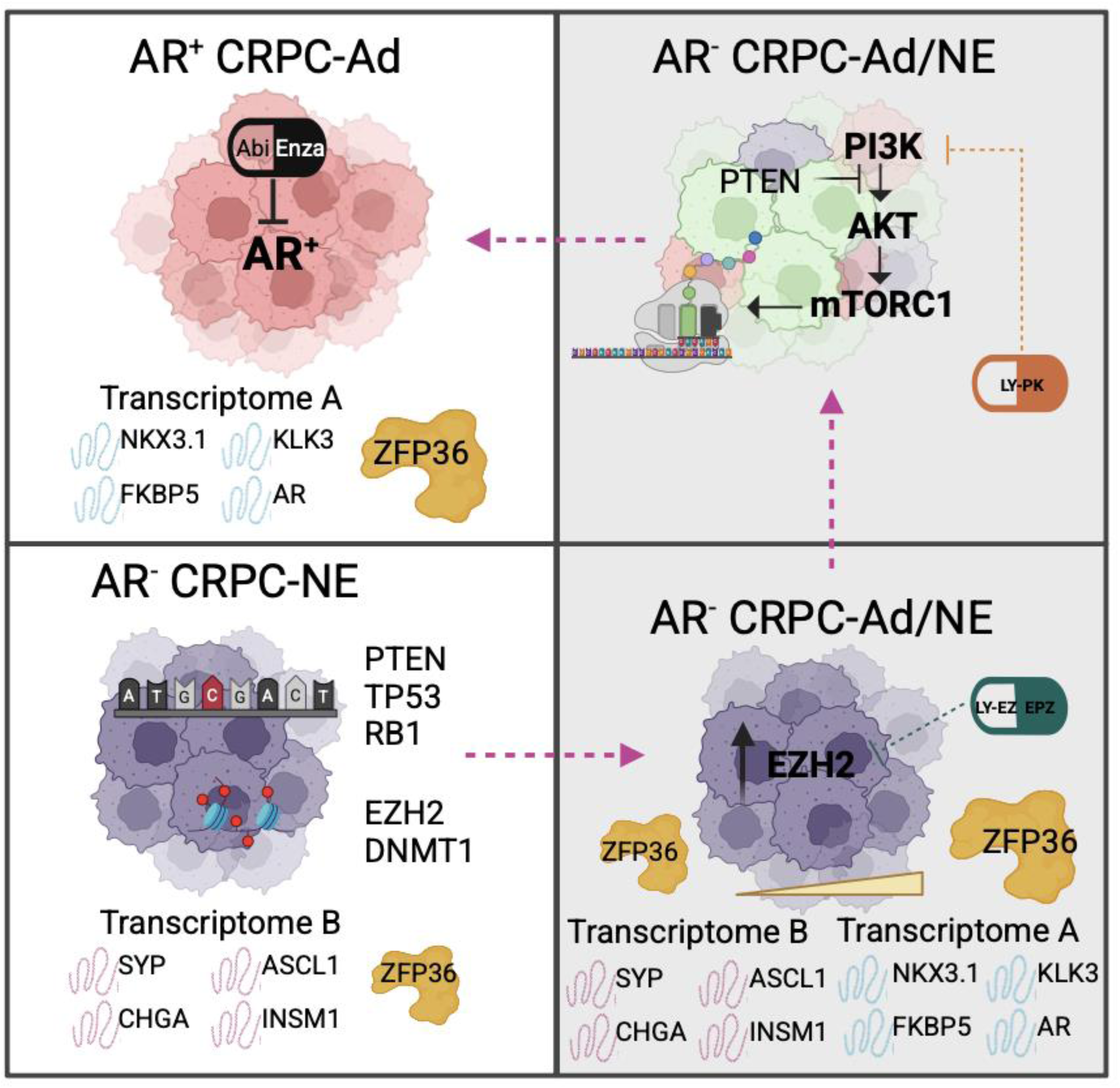

**Significance Statement:** EZH2 plays a key role in driving cell state transitions in neuroendocrine prostate cancer (NEPC), guiding cancer cells towards a more aggressive, therapy-resistant cell type. This transformation creates a specific vulnerability, as NEPC cells become highly reliant on both EZH2 and PI3K/mTORC1-translation signaling. Targeting this dependency, we demonstrate that combining EZH2 with PI3K/mTORC1 inhibition provides effective suppression of NEPC cell growth, offering a promising therapeutic strategy for treating this challenging and aggressive prostate cancer subtype.

## Introduction

Targeting androgen biosynthesis and androgen receptor (AR) function remains the cornerstone of treatment for prostate cancer (PCa) patients who have progressed beyond surgery and/or radiation therapies. While androgen receptor signaling inhibitors (ARSIs) offer a survival benefit, resistance to these therapies is often inevitable. Primary mechanisms driving resistance to ARSI involve the restoration of AR expression and function via AR amplification, mutation, or the expression of constitutively active AR splice variants, all of which sustain the AR-driven transcriptional program. Additional evidence now demonstrates that approximately 15-20% of patients develop ARSI resistance through a distinct mechanism termed phenotypic plasticity. This adaptive process involves epigenetic reprogramming and transcriptomic remodeling, often driven by loss-of-function alterations in key tumor suppressors such as PTEN, RB1, and TP53, which decouple tumor growth from AR dependency (1–3).

These PCa often display altered kinase signaling and multilineage states involving activation of alternate transcriptomes (4) driving divergent cell identities including neuroendocrine-, stem-, and basal-like gene signatures (2, 3, 5, 6). These lineage-divergent states are marked by the emergence of alternate transcriptomes and lineage master regulators, contributing to therapeutic resistance and tumor aggressiveness. Importantly, phenotypic plasticity in PCa parallels similar processes observed in other solid tumors, including *EGFR*-mutant lung adenocarcinoma, Merkel cell carcinoma, and *BRAF* mutant melanoma, where transdifferentiation also underlies therapeutic resistance (7–9). Although increasing insights into the molecular drivers of this plasticity, successful translation into effective therapies remains elusive. Research, including our own, has shown that a prominent mediator of plasticity is EZH2, the catalytic subunit of the Polycomb Repressive Complex 2 (PRC2), which represses gene expression through H3K27 trimethylation (10). In tumors exhibiting phenotypic plasticity, EZH2 is frequently overexpressed and contributes to silencing of differentiation-associated genes, thereby sustaining alternate lineage states. In this context, loss of PRC2 in mouse models leads to the emergence of bivalent promoters marked by both active H3K4me and repressive H3K27me3 marks, features typically acquired during differentiation (11–14). Beyond its canonical methyltransferase activity, EZH2 also exerts non-catalytic functions, such as modulating mRNA translation and interacting with components of the translational machinery, that are recognized as relevant to tumor biology, including the role of EZH2 inhibition restoring AR expression, and response to enzalutamide (2, 3, 15).

Further, aberrant activation of kinase signaling pathways has been reported in metastatic castration resistant PCa (mCRPC) and is particularly pronounced in tumors undergoing phenotypic plasticity (16, 17). Among these, the Janus kinase (JAK)-signal transducer and activation of transcription (STAT) signaling has emerged as key driver of phenotypic plasticity, and inhibition of JAK-STAT signaling re-sensitized phenotypic plastic PCa to enzalutamide via restoration of luminal lineage identity and AR signaling (5, 18). These findings emphasize the importance of inflammatory and cytokine-driven signaling networks in facilitating lineage reprograming.

The PTEN/AKT/mTOR pathway, frequently activated through PTEN loss, is another critical axis in PCa, promoting cell survival, proliferation, and resistance to therapy. Canonically, EZH2 suppression inhibits mTORC1 signaling by derepressing its negative regulators such as TSC2, PTEN, and DEPTOR, leading to reduced phosphorylation of mTORC1 downstream effectors, e.g., 4E-BP1 and S6K, and subsequent induction of autophagy and apoptosis (19–22). EZH2 can also repress translation through direct interaction with eIF4E, independently of histone methylation (23). However, in specific oncogenic contexts such as NrasG12D-driven acute myeloid leukemia, EZH2 loss paradoxically activates mTORC1 through increase of intracellular branched-chain amino acids and mTORC1 activation (24). These context-specific findings underscore the dual, and sometimes opposing, regulatory effects of EZH2 on mTOR signaling, protein synthesis, and cancer progression.

Here, using a genetically engineered mouse model (GEMM) of phenotypic plasticity driven by concurrent *Pten*:*Rb1* deficiency, we demonstrate that EZH2 regulates multilineage states and their master-regulator transcription factors (MR-TF). Inhibition of EZH2 partially reverses these cell states and displays dependence on translational activation, ribosomal biogenesis, and RNA stability. These molecular shifts revealed a vulnerability that could be therapeutically exploited. Co-inhibition of EZH2 and the PI3K/AKT/mTORC1 pathway produced superior antitumor activity in both murine and human preclinical models of plastic PCa, especially in castration settings where AR signaling is suppressed. Together, these data position EZH2 as a key epigenetic gatekeeper of phenotypic plasticity in PCa and therapeutic resistance. Targeting EZH2 in combination with translational and kinase signaling inhibitors may thus offer a powerful strategy to overcome lineage plasticity, restore treatment sensitivity, and suppress progression in aggressive PCa.

## Results

### EZH2 coordinates multi-lineage cell states mediated by RB1 loss

We previously demonstrated that co-loss of *Pten* and *Rb1* (DKO mice (3)) genes promotes a shift from prostate adenocarcinoma to a NE-like PCa phenotype that is resistant to ARSI. A key consequence of *Rb1* loss is the upregulation of EZH2 (3), which mediates widespread chromatin remodeling, including the establishment of bivalent chromatin domains marked by both H3K27me3, associated with gene repression, and H3K4me3, associated with gene activation (11, 13). To explore this epigenetic reprogramming, we performed immunoprecipitation following by sequencing (ChIP-seq) for the histone modifications H3K4me3 and H3K27me3 in DKO (NE-like model) and SKO (*Pten* KO, adenocarcinoma-like model) GEMMs. This analysis revealed a global chromatin redistribution of these marks in DKO tumors (**Fig. S1A-D**). In particular, we observed a notable repositioning and enrichment of H3K27me3 (**Fig. S1B)**, supporting the model in which EZH2 activity drives chromatin remodeling and transcriptional repression of lineage-determining genes. Further, when EZH2 was inhibited, partial reversal of the NE-like cell state was achieved and re-sensitized tumors to AR antagonism (3). Here, DKO GEMMs were treated at approximately 38 weeks with either control (n=2) or the EZH2 inhibitor, EPZ011989 (n=2) for 1 week. The tumors were then subjected to single nuclei RNA sequencing (snRNA-seq). A total of 10,000 *Epcam*^+^ (epithelial) cells were analyzed using an unbiased Leiden method and visualized by uniformed manifold approximation and projection (UMAP) (**Fig. 1A**). UMAP distribution based on treatment revealed that EZH2 inhibition was responsible for 9 out of the 10 Leiden clusters originally generated (**Fig. 1A**), implicating EZH2 as an important mediator of multilineage cell states by controlling alternate transcriptomes. These results are in line with recently reported data from mouse and human PCa devoid of *RB1* and/or *TP53* function (5).

**Figure 1:**
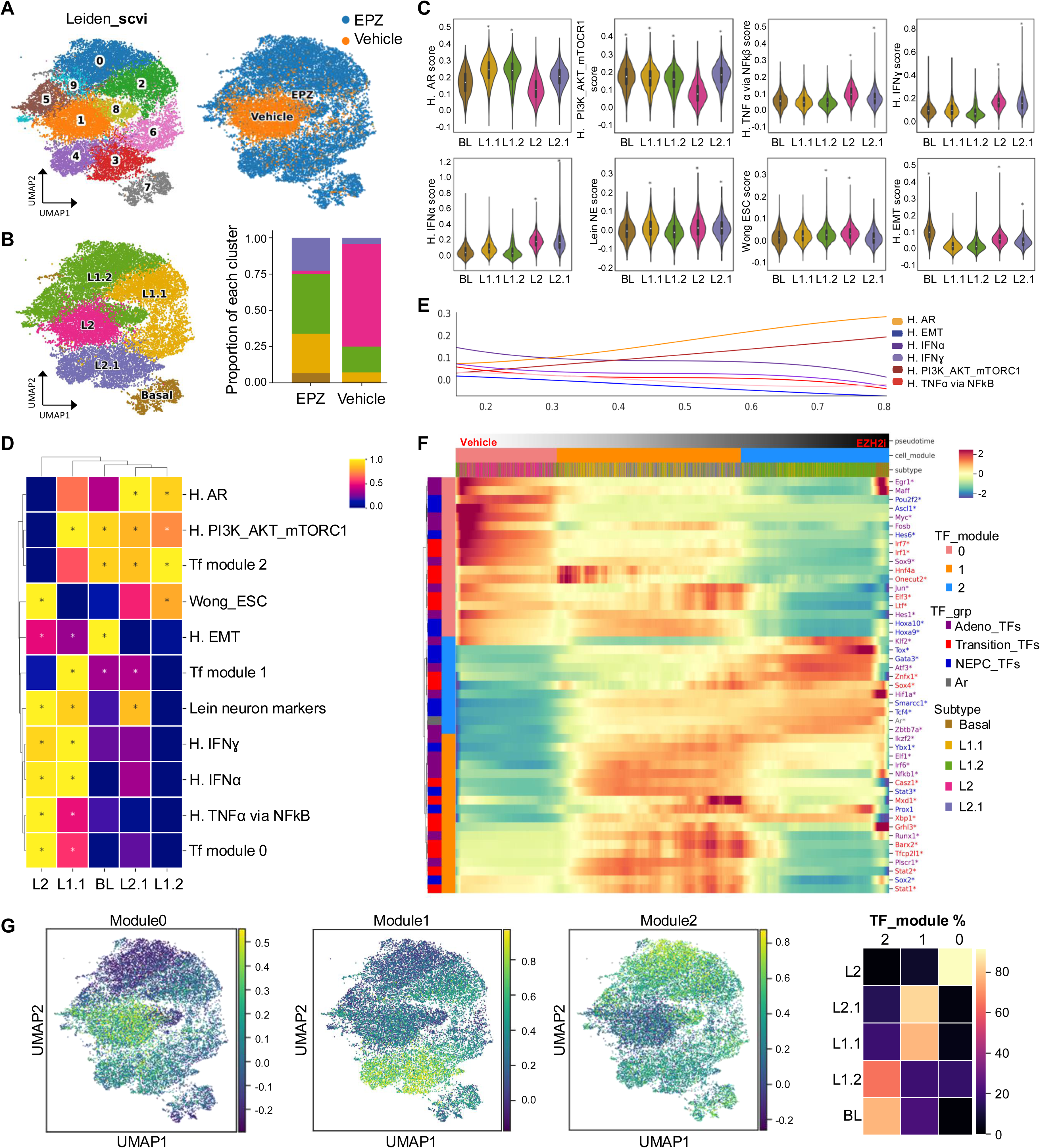
EZH2 coordinates multi-lineage cell states downstream of RB1 loss. *PbCre:Pten:Rb1* KO (DKO) mice were treated daily for 7 days with vehicle (n=2) or EZH2 inhibitor EPZ011989 (EZH2i, n=2) via oral gavage at 38 weeks of age. Tumors were harvested, dissociated, and processed for single-nucleus RNA sequencing (snRNAseq). **A**. Left panel represents the Uniform Manifold Approximation and Projection (UMAP) visualization of Leiden-based unsupervised clustering of the epithelial cells (Epcam^+^). Right panel represents the UMAP showing treatment specific distribution of epithelial cells from EZH2i-treated (blue) and vehicle-treated (orange) tumors. **B.** UMAP and bars plot depicting epithelial subtype classification including basal-like (BL), luminal 2 (L2), luminal 2.1 (L2.1), luminal 1.1 (L1.1), and luminal 1.2 (L1.2) cells. **C-D.** Violin plots and heatmaps showing significantly enriched gene sets across epithelial subtypes (BL, L1.1, L1.2, L2 and L2.2) in EZH2i-treated DKO tumors. Gene set enrichment analysis (GSEA) was performed with FDR < 0.01 and absolute normalized enrichment score (NES) > 1 (see methods). H. AR: hallmark_AR (NELSON_ANDROGEN_SIGNALING_UP), H. PI3K_AKT_mTORC1: hallmark_ PI3K_AKT_mTORC1, H. IFNɣ: hallmark interferon gamma, H. TNFα signaling via NFkB: hallmark TNFα signaling via NFkB, H. IFNα: hallmark of interferon alpha, Wong ESC: Wong embryonic stem cells, H. EMT: hallmark of epithelial mesenchymal transition. **E.** Pseudotime analysis representing a spline-fitted average Z-scores of hallmark gene sets (hallmark of AR signaling, EMT, IFNα, IFNɣ, PI3K-AKT-MTOR, and TNFα signaling via NFkB) (see methods) from vehicle to EZH2i-treated mice. **F.** Heatmap of master regulatory transcription factors (MR_TFs) implicated in the transition from prostate adenocarcinoma to neuroendocrine prostate cancer (NEPC) across Epcam^+^ cells. Transcription factors groups (TF_grp) include adenocarcinoma-associated (adeno_TFs, purple), transition-associated (transition_TFs, red), and NEPC-associated (NEPC_TFs, blue). Cells are categorized into TF modules (modules 0-2) based on pseudotime, epithelial subtype, and MR-TF expression (scale imputed expression range −0.5 to 1.5). **G.** UMAP and heatmap visualization of the TF_grp expression across the TF_modules.

We further defined the sub-clusters using specific gene-sets previously published which identify cell types during prostate development and PCa progression (5, 25–27) (**Fig. 1B** and **Fig. S2A**). Additional interrogation using Gene Set Enrichment Analysis (GSEA) allowed visualization of specific pathways both unique and shared between sub-clusters (**Fig. 1C-D, Fig. S2B-C** and **Table S1**). The DKO (vehicle treated) cluster defined as L2 (luminal 2) demonstrated most multilineage cell states represented by enrichment of gene signatures involving inflammation, epithelial-mesenchymal transition (EMT), neuron markers, and stemness, and exhibited most repression of AR genes and PI3K/AKT/mTORC1 signaling (**Fig. 1C-D** and **Fig. S2B-D**). These gene signatures were notably inversed with EZH2 inhibition, which generated sub-clusters L2.1 through to L1.2 suggesting that EZH2 tightly regulates control of gene expression and transition of these noted epithelial cell states (**Fig. 1C-D, Fig. S2D**). A final cell state that was generated by EZH2 inhibition was identified as a basal-like cell population (**Fig. 1B-D**). While this cluster exhibited strong basal markers (**Fig. S2A**), it also exhibited strong enrichment of EMT, PI3K/AKT/mTORC1, and moderate AR gene signatures (**Fig. S2A**). Utilizing similar GEMMs, recent work performed single cell RNA sequencing (scRNA-seq) to investigate causes of phenotypic plasticity in PCa (5). From this scRNA-seq analysis a list of master regulator transcription factors (MR-TF) was generated and defined as i) those active in adenocarcinoma, ii) those active in a putative transition to NEPC, and iii) those specific of NEPC induction (5). Applying this MR-TF list to our snRNA-seq data we observed NEPC related MT-TFs were enriched within our DKO vehicle treated (L2) population and EZH2 inhibition generated subpopulations enriched for transitional and adenocarcinoma MR-TFs (**Fig. 1E-F**). We confirmed these observations by visualizing these MR-TF modules on our UMAP (**Fig. 1G**). We specifically noted spatial distribution with NEPC MR-TFs enriched within L2 and L2.1 subpopulations (module 0), transition MR-TFs enriched in L2.1, basal, and L1.1 subpopulations (module 1), and adenocarcinoma MR-TFs enriched in basal, L1.1, and L1.2 (module 2), again emphasizing the important control by EZH2 on these cell state transitions (**Fig. 1F-G**). To investigate how EZH2 shapes the epigenetic landscape and contributes to phenotypic plasticity in PCa, based on the ChIP-seq profiles of H3K4me3 and H3K27me3 we classified the chromatin into four major states: (i) Active (H3K4me3+ only), (ii) Repressed (H3K27me3+ only), (iii) Bivalent (both H3K4me3+ and H3K27me3+), and (iv) Unmarked (neither mark). This classification enabled us to track chromatin state transitions from SKO to DKO tumors and to identify chromatin states and lineage-specific genes most targeted by EZH2 inhibition (**Table S2**). To do so, we integrated our ChIP-seq data with the snRNA-seq analysis from DKO tumors treated with an EZH2 inhibitor. Notably, we observed that bivalent chromatin states in DKO tumors were most significantly targeted by EZH2 inhibition (**Fig. S1F**), suggesting a re-establishment of developmental poising. Examples of this remodeling involved in this transition include Notch1, Tacstd2, or Mycn which are up regulated in DKO tumors upon EZH2 inhibition (**Fig. S1E** and **Table S2**). These findings indicate that EZH2 inhibition induces broad epigenetic reprogramming, including redistribution of H3K27me3, which is accompanied by alterations in epithelial cell states. Together, our data highlight a critical role for EZH2 in maintaining phenotypic plasticity in prostate cancer models involving *Rb1* deletion and suggest that its inhibition may restore chromatin dynamics that constrain lineage infidelity.

### Canonical and non-canonical function of EZH2 identifies translation as a dependency for phenotypic plasticity

To further identify dependencies that could serve as potential therapeutic targets, we first employed a functional genomics approach. DKO cells expressing Cas9 nuclease were transduced with the Brie lentiviral gRNA library, which contains 78,637 gRNAs targeting 19,674 protein-coding genes in the mouse genome (28). Following several days of treatment with either DMSO or the EZH2 inhibitor, EPZ6438, genomic DNA was extracted, and sgRNA abundance was quantified using high-throughput sequencing (**Fig. 2A**). The data were analyzed using the MAGeCK algorithm (29, 30) (**Table S3**). Analysis, of depleted genes revealed significant enrichment in gene ontology (GO) terms related to translation, RNA processing, and ribosome biogenesis (**Fig. 2B**). In parallel, we conducted an independent proteomic analysis to assess the potential differential role of EZH2 as a transcriptional co-factor. Using SKO and DKO cells, we performed rapid immunoprecipitation mass spectrometry of endogenous peptides (RIME) (31) (**Fig. 2A**) following EZH2 protein pulldown. To nominate differential EZH2 binding partners, we applied a false discovery rate (FDR) cutoff of 0.1, requiring enrichment relative to the IgG control and detection in at least two out of the four samples **(Table S4**). GO term enrichment of RIME-identified interactors mirrored the CRISPR screen results, with significant representation in translation, RNA processing, and ribosome biogenesis pathways. Interestingly, GSEA using the Hallmark gene sets further demonstrated that both datasets converged on mTORC1 signaling (**Fig. S3**), consistent with results from our snRNAseq analysis (**Fig. 1, Fig. S2**). To validate the modulation of mTOR signaling by EZH2 inhibition, we performed intracellular flow cytometry in DKO cells treated with EZH2 inhibition, assessing levels of phospho-4E-BP1 (p-4E-BP1), phospho-S6K (p-S6K) and TSC2. EZH2 inhibition led to a significant 3-fold increase in p4E-BP1 levels relative to DMSO control (p<0.001), consistent with enhanced translation initiation (**Fig. 2C**). In contrast, p-S6K levels remained unchanged (fold change around 1; p>0.05), suggesting that this branch of the mTORC1 pathway is not affected under these conditions (**Fig. 2C**). Notably, EZH2 inhibition also resulted in a significant increase in TSC2 protein expression (fold change relative to control; p=0.012) (**Fig. 2C**). The concurrent upregulation of p4E-BP1 and TSC2 suggests a global activation of translation programs rather than a direct regulatory interaction between the two proteins.

**Figure 2:**
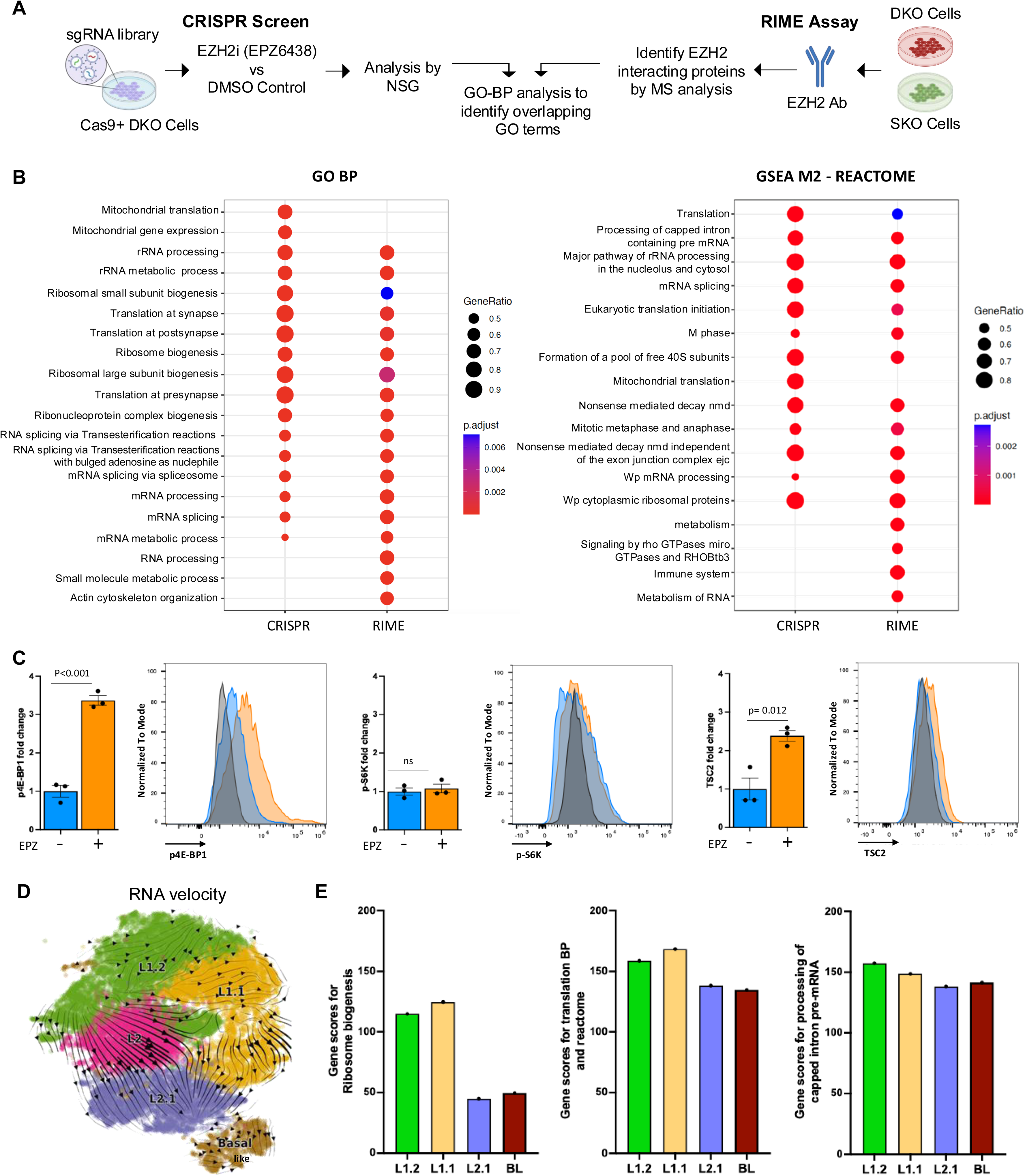
EZH2 Inhibition Identifies Coordination of Translation as a Genetic Dependency in Rb1 Deficient Prostate Cancer. **A.** Schematic overview of two complementary approaches to identify EZH2 regulators and interactors. (Left) In the positive selection CRISPR screen, DKO cells were transduced with the Brie lentiviral gRNA library that consists of 78,637 gRNAs that target 19,674 protein coding genes and subjected to EZH2i (EPZ6438) treatment. Surviving/enriched populations were sequenced to identify genes whose loss conferred resistance or enhanced survival. (Right) In parallel, Rapid Immunoprecipitation Mas spectrometry of endogenous proteins (RIME) was performed in SKO (Pten KO) and DKO (Pten:Rb1) cells using anti-EZH2 antibody to immunoprecipitated endogenous protein complexes from crosslinked chromatin. Mass spectrometry analysis identified associated proteins, revealing potential co-regulators and chromatin–bound interactors**. B.** Dot plot showing gene ontology biological process (GO-BP) and Reactome pathway enrichment across hits from the CRIPR screening and RIME analysis in SKO and DKO cells. Color indicates the p-value, and dot size reflects the percentage of cells expressing the pathway-associated genes. **C.** Bar graph and histogram showing fold changes in TSC2, phosphorylated 4E-BP1 (p4E-BP1), and phosphorylated S6K (p-S6K) levels in DKO cells treated with EZH2 inhibitor (EPZ6438) versus DKO cells treated with DMSO. Data represent mean ± SEM from [n=3] independent experiments. EZH2i treatment significantly increased TSC2 and p4E-BP1, while p-S6K remained unchanged, indicating a selective modulation of mTORC1 signaling. ns= no significant. **D.** RNA velocity analysis of epithelial cells in vehicle and EPZ treated mice. **E.** Bar plots showing the signature related with translation mechanisms in L2 sub-cluster versus other sub-clusters (L2.1, L1.1, L1.2 and BL). The analyzed signatures include genes related with ribosome biogenesis, translation at synapse, post-synapse and pre-synapse, and translation (in biological processes and reactome). The comparison highlights differential expression of key translational regulators in response to EPZ treatment across cellular subtypes.

To assess the relevance of translational programs under EZH2 inhibition, we examined gene score signatures related to translation and ribosome biogenesis, identified via CRISPR and RIME analysis, within the differentially expressed gene profiles of sub-clusters derived from EZH2i-treated mice (L1.2, L1.1, L2.1, and BL) versus vehicle-treated (L2) cluster. RNA velocity analysis, which infers future cell states and transitions across subclusters (32), indicated that EZH2 inhibition drives cell fate progression from the L2 (DKO vehicle treated cells) cluster toward the ‘most differentiated – L1.2’ and to ‘least differentiated – L2.1 cell states’, eventually leading to a basal-like cell population (**Fig. 2D**). Pseudotime analysis confirmed this trajectory (33) (**Fig. S3B**). Notably, ribosome biogenesis gene sets were enriched primarily in L1.2 ad L1.1 clusters, while translation-related and processing of capped intron pre-mRNA signatures were uniformly distributed across subclusters. This pattern paralleled a consistent increase in PI3K/AKT/mTORC1 signaling score (**Fig 2E and Fig S2D and S3D**). These observations implicate that EZH2 mediates phenotypic plasticity at multiple levels involving canonical catalytic functions and potentially non-catalytic functions as part of the transcriptional complex.

### Combination of EZH2 and PI3K/mTORC1 inhibition as a novel therapeutic approach in phenotypic plastic prostate cancer

From our data, we have demonstrated that EZH2 inhibition enriches for mTORC1 signaling, and our functional genomics and proteomics data further implicate EZH2 to be significantly involved with translational control in models of phenotypic plasticity. Previously it has been demonstrated that aberrant translation in PCa tumoral cells is required to maintain AR-low PCa cells promoting tumor heterogeneity (34). Therefore, we proposed that combinatorial inhibition of EZH2 with mTORC1 inhibition would provide superior anti-tumor efficacy. Treatment of DKO (*Pten:Rb1* KO) and PPKO (*Pten:Trp53* KO) spheroids with two independent EZH2 inhibitors, LY3346149 (LY-EZ), and EPZ6438, in combination with LY3023414 (LY-PK, a dual PI3K/mTORC1/2/DNA-PK inhibitor) for 3 days, showed that castrate (CSS) conditions presented the most significant synergy compared to androgen replete (FBS) conditions (**Fig. 3A-B, Table S5,** and **Fig. S4A**). Next, we investigated DKO response in vivo by injecting intact, or castrated mice with DKO cells. These mice were treated with vehicle control, LY3346149 (LY-EZ), LY3023414 (35) (LY-PK), or combination (**Fig. 3C**). Specific to both intact and castrated treated groups, EZH2 inhibition alone did not provide anti-tumor efficacy, whereas response to PI3K/mTORC1 inhibition alone provided significant anti-tumor effect against intact tumors, but acquired resistance was noted in castrated conditions (**Fig. 3D)**. No observed advantage in anti-tumor effect occurred when combination treatment was compared to LY-PK monotherapy of intact DKO tumors. The resistance noted to LY-PK monotherapy was reversed by the addition of EZH2 inhibition, suggesting the acquired resistance seen with DKO castrate-resistant tumors is mediated by EZH2 catalytic function (**Fig. 3D**). The analysis of formalin-fixed paraffin embedded (FFPE) tissue showed loss of H3K27me3 in both intact or castrated tumors at comparable levels when treated with EZH2 inhibition or combination. Further, combination therapy also generated significant loss of overall tumor cell proliferation (Ki67), and increased double-strand DNA damage as measured by p-γH2AX staining (**Fig. S4B-C**). Importantly, using a second independent murine model with co-deletion of PTEN and TP53 (36) (PPKO), and a human NEPC organoid model (37) when transplanted in vivo demonstrated similar results to what was observed in our DKO in vivo model (**Fig. 3E-F**). Both PPKO and human NEPC models displayed modest response when treated with the single agent EZH2 or PI3K/mTORC1. Like DKO results, EZH2 and PI3K/mTORC1 inhibitor combination plus castration or enzalutamide provided most significant anti-tumor activity. We had previously demonstrated that EZH2 inhibition, by EPZ6438, in DKO cells could restore AR expression and function explaining the synergy with castration or enzalutamide (3). This held true with the LY-EZ compound demonstrating robust increase expression and function of AR in both DKO and PPKO cells (**Fig. S5**).

**Figure 3:**
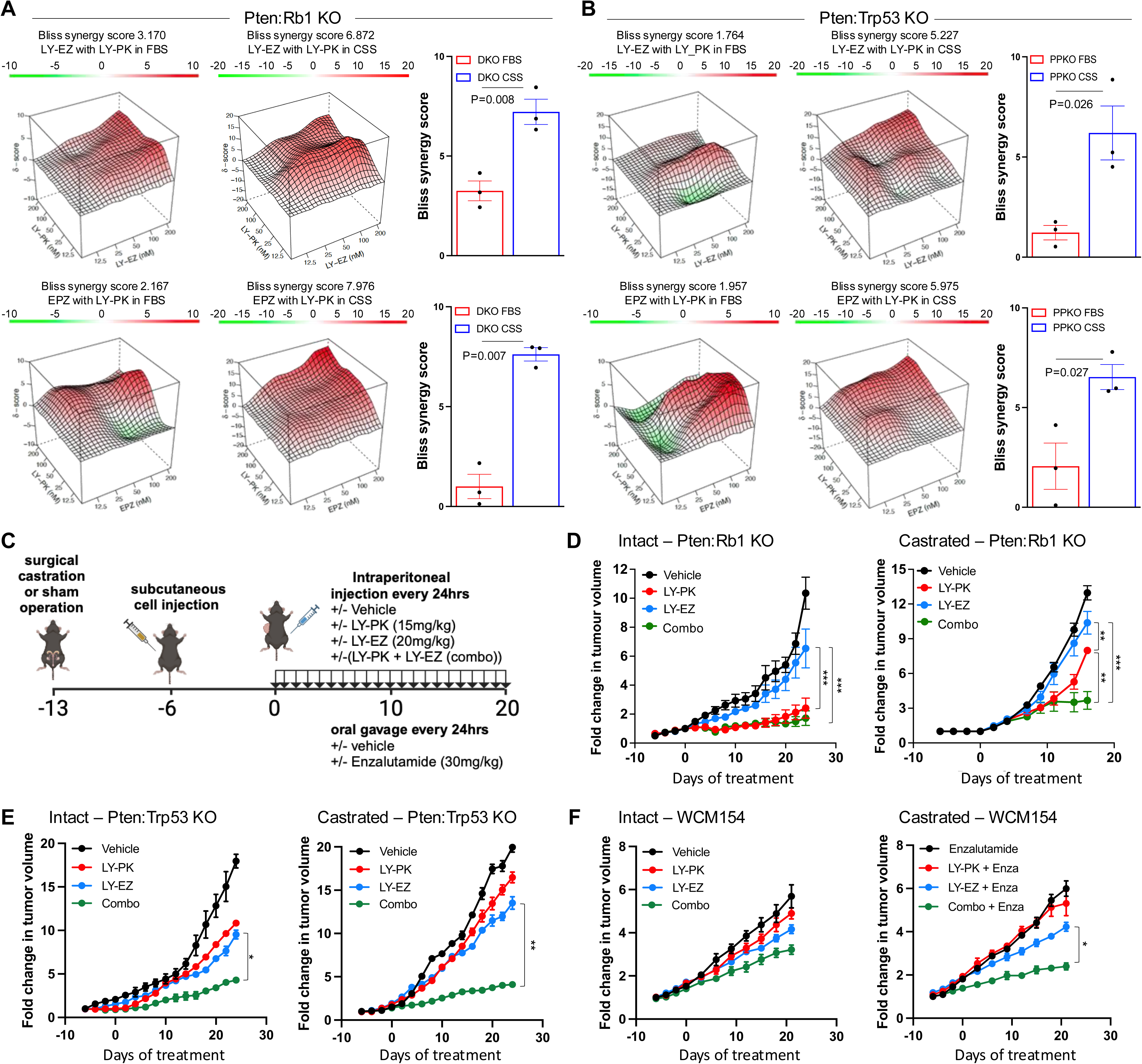
Combination of EZH2 and PI3K/mTORC1 inhibition as a novel therapeutic approach in phenotypic plastic prostate cancer. **A-B.** Bliss synergy landscapes generated using the Bioconductor package *synergyfinder* for drug combinations in 3D spheroid models. Graphs represent one representative replicate (with maximal synergy closest to its average value). Synergy score (z-axis) ranges from −20 (green, antagonism) to +20 (red, strong synergy). X- and Y- axes represent the concentrations of the two drugs. A. Drug combinations (LY-EZ with LY-PK (top), and EPZ with LY-PK (bottom)) tested in DKO spheroids under FBS and CSS (charcoal-stripped serum) conditions. **B.** Same combination tested in *Pten:Trp53* KO (PPKO) spheroids. For specific drug concentrations, refer to Table S6. **C.** Schematic representation of the experimental procedure to evaluate the therapeutic effect of combinatory therapies in three independent preclinical models. D-F. Tumor growth curves showing fold change in tumor size over time under different treatment conditions (LY-PK, LY-EZ, and the combination). **D.** Graphs representing the growth curves for DKO GEMM-derived allografts in C57BL/6N mice in intact or castrated conditions upon the specific treatments. In brief, C57BL/6N mice were surgical castrated, or sham operated at 8 weeks of age, and after 7 days castrated resistant 1×10^6^ DKO cells were implanted into the flank of each mouse in 100μL of Matrigel (50% in PBS). Once the tumors acquired 20mm^2^ mice were treated daily with LY-PK (15 mg/kg) and/or LY-EZ (20 mg/kg) by intraperitoneal (IP) injection. During treatment tumors were measured every second day for 20-30 days (n=5 mice per treatment group (± 1SEM). **E.** Graphs representing the growth curves for PPKO GEMM-derived allografts in C57BL/6N mice in intact or castrated conditions upon specific treatments. **F.** Graphs representing the tumor growth of 1×10^7^ human neuroendocrine PCa model (WCM154) in NOD.CB17-Prkdcscid/NCrCrl (NOD/LtSz) mice in intact or castrated conditions upon different treatments, following same schema than in C.

### The RNA binding protein Tristetraprolin (TTP) is a dependency for EZH2 mediated cell state transition and therapeutic efficacy

Our results demonstrate that EZH2 inhibition resulted in the enrichment of translation signaling pathways and the de-enrichment of inflammatory pathways mediated by interferon alpha (IFNα), interferon gamma (IFNγ), and TNFα-NKκB signaling. We have previously demonstrated using a *Pten* KO GEMM of PCa, that co-deletion of TTP (*ZFP36,* gene which encodes for TTP) resulted in significant changes in gene expression, predominantly inflammation and cell identity related gene-sets (38). Our findings led us to investigate whether TTP function plays a role in the cell state transitions mediated by EZH2 and overall therapeutic response. Previously published gene expression data from mouse (3) and human (2) PCa samples revealed that *ZFP36* is significantly downregulated in phenotypic plastic/NEPC compared to adenocarcinoma samples (**Fig. 4A**). This was also determined at protein level in our murine cell lines (**Fig. 4B**). Moreover, treatment of DKO cells with an EZH2 inhibitor resulted in an increase in TTP expression (**Fig. 4B**). This regulation of *ZFP36*/TTP expression and function by EZH2 was also observed in our snRNAseq data where application of *ZFP36* gene signature derived from our *Pten*:*Zfp36* KO GEMM (38) showed that EZH2 inhibition resulted in downregulation of the gene signature, indicating that TTP function was activated (**Fig. 4C**). Because of *ZFP36*/TTP activation by EZH2 inhibition, we developed *ZFP36* knockout DKO and PPKO cells (**Fig. S6A**). Treatment of the DKO-*Zfp36* KO cell line with two independent EZH2 inhibitors, LY-EZ and EPZ6438, followed by RNA sequencing revealed that cells depleted of *Zfp36*/TTP were unable to downregulate inflammatory gene sets and significantly increase AR and mTORC1 related gene signatures (**Fig. 4D**) as previously demonstrated (**Fig. S2B**). The dependency of *Zfp36/*TTP shown for EZH2 inhibition was further demonstrated when combination treatment of EZH2 with PI3K/mTORC1 inhibition was not efficient to control the cell growth of the DKO-*Zfp36* KO and PPKO-*Zfp36* KO models (**Fig. 4E-F** and **Table S6**).

**Figure 4:**
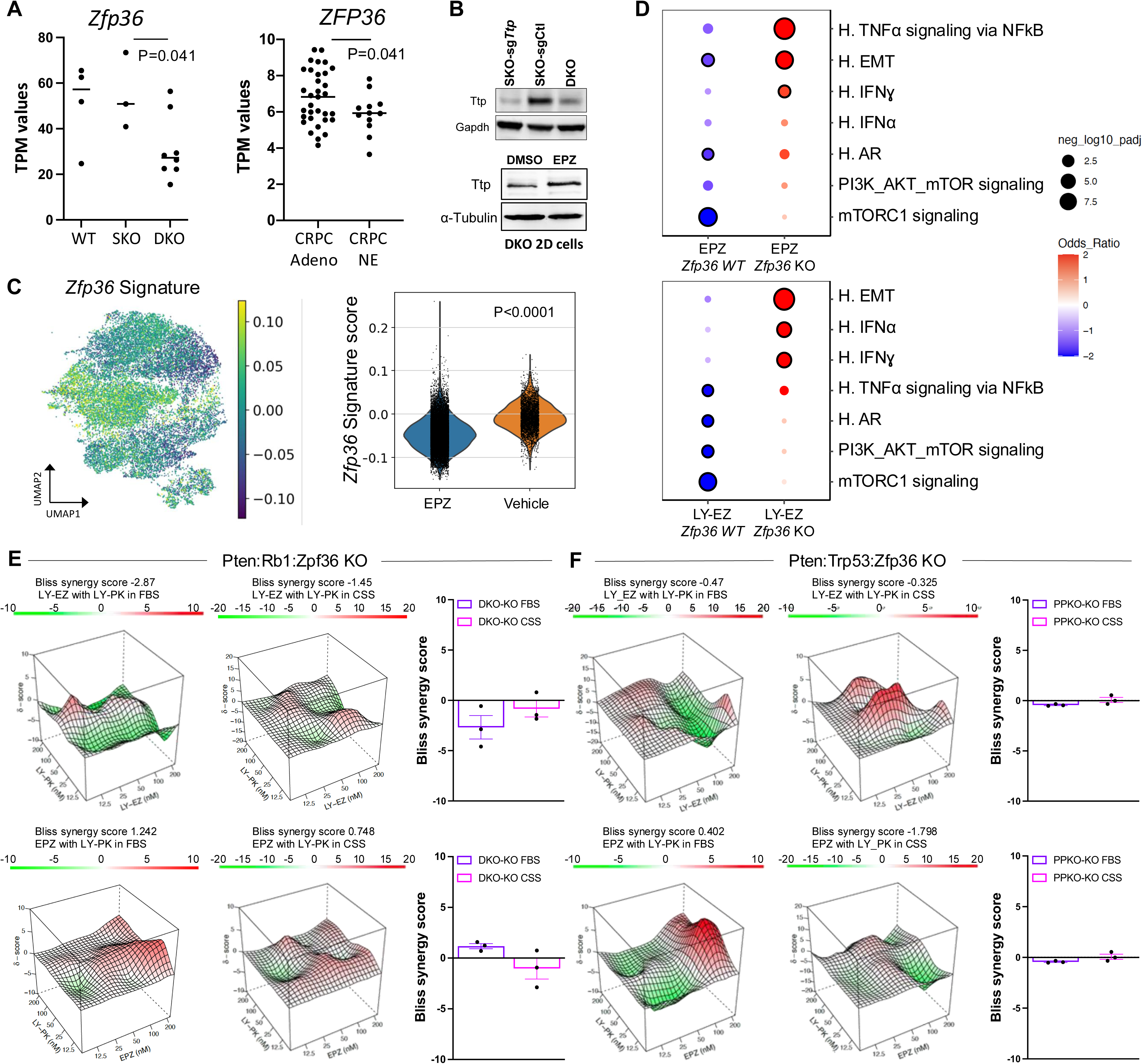
EZH2 Inhibition Cell State Transition Dependent on the RNA Binding Protein Tristetraprolin (TTP) **A.** RNAseq analysis of GEMM (wild type (WT), SKO, DKO) and human (castration resistant prostate cancer (CRPC) and Neuroendocrine prostate cancer (NEPC)) samples showing consistent downregulation of Zfp36/ZFP36 in NEPC. **B.** Western blot showing Ttp protein levels in SKO Zfp36 KO, SKO control (Ctl), and DKO control cells (top) and DKO cells treated with DMSO or EPZ (5uM) (bottom). **C.** UMAP and violin plot of the *Zfp36* gene target signature generated from a our *Pten:Zfp36* KO GEMM demonstrating EZH2 induces Ttp (Zfp36) function. **D.** Dot plot of GSEA comparing gene signatures upregulated in EPZ or LY-EZ treated DKO *Zfp36* WT and KO cells. Color intensity represents average normalized enrichment score; dot size indicates the percentage of expressing cells. **E-F.** Bliss synergy landscapes for LY-PK combined with LY-EZ or EPZ6438 in *Zfp36* KO spheroids. For specific drug concentrations, refer to Table S6. E. Examples of the synergy profiles in DKO-*Zfp36* KO spheroids under FBS and CSS conditions. F. Example of the synergy profiles in PPKO-*Zfp36* KO spheroids under the same conditions.

## Discussion

Phenotypic plasticity has emerged as a new hallmark of cancer and a growing concern as a mechanism driving therapeutic resistance. Specifically in PCa, phenotypic plasticity is thought to be a gateway for prostate adenocarcinoma to evolve by induction of a cell state switch to where that cell harbors a new molecular identity often referred to as neuroendocrine PCa (NEPC) (3, 5, 39–41). Previously we observed that EZH2 inhibition partially reversed this lineage shift and restored sensitivity to ADT in part by restoration of AR expression and function (3). To more deeply address the regulation of cellular changes governed by EZH2, we performed snRNA-seq using a PCa GEMM of phenotypic plasticity/NEPC driven by concurrent loss of *Pten* and *Rb1* (DKO).

Treating DKO with control or with a EZH2 inhibitor and using previously published PCa scRNAseq datasets (5, 34, 42, 43) to annotate cell identity, we found that vehicle treated DKO tumors predominately consist of L2/L2.1 progenitor like cells (*Psca*^+^/*Krt4^+^/Tacstd2^+^*) with smaller subsets of L1/L1.1/L1.2 intermediate/differentiated luminal cell types, and luminal-basal cell populations. Following EZH2 inhibition, the L2 subpopulation was drastically decreased, while L2.1, L1 related subpopulations, and luminal-basal cells were dramatically increased. GSEA across subgroups further supported previous data (3, 5) where gene sets related to interferon and NFκB inflammatory signaling, neuronal markers, and stemness were strongly enriched in DKO cells. Inhibition of EZH2 controlled depression of these gene-sets and an increased enrichment of AR signaling, that we previously reported (3), and PI3K-AKT-mTORC1 signaling were observed. Interestingly, while EMT genes were enriched in L2 subpopulations, they were most increased in the luminal-basal cell population. These data coincide with previously publications demonstrating the importance of basal and basal-luminal cells for the initiation of PCa and phenotypic plasticity (5, 44–47). In addition, MR-TFs driving transition of prostate adenocarcinoma to a NEPC phenotype were recently described (5). These MR-TFs were defined into 3 main modules representing adenocarcinoma (module 2), transition (module 1), and NEPC (module 0). These MR-TF modules were clearly represented with the NEPC module MR-TFs enriched in DKO-L2 (vehicle treated), and the transition and adenocarcinoma module MR-TFs enriched in DKO-L2.1 and DKO-L1 subgroups respectively. Collectively, our data clearly demonstrate the importance for EZH2 catalytic function to govern expression/action of MR-TFs that drive phenotypic plasticity and cell state transitions allowing prostate adenocarcinoma evolution to NEPC to evade AR signaling inhibitors.

While EZH2 is well established as a histone methyltransferase and transcriptional repressor within the PRC2 complex, accumulating evidence suggests that it also exerts non-canonical functions independent of its methyltransferase activity. These include roles in transcriptional activation (48, 49), interaction with transcriptional machinery (50), and modulation of oncogenic signaling pathways such as AKT and Wnt (51, 52). Building on this evolving understanding, our findings suggest that EZH2 may also influence translational regulation. Through integrated functional genomic and proteomic approaches, we observed a genetic dependence on translational-related processes in EZH2-driven contexts. Notably, EZH2-interacting proteins were enriched in pathways related to ribosome biogenesis and mRNA processing. We further observed elevated p4E-BP1 and increased TSC2 protein levels, accompanied by stable p-S6K expression, indicating a selective modulation of the mTORC1 axis, potentially favoring initiation of cap-dependent translation. These results are consistent with prior reports suggesting that translational reprograming can contribute to phenotypic plasticity in cancer (49, 53). Trajectory analysis of snRNAseq further revealed that EZH2 inhibition was associated with transitions across distinct states, accompanied by increased PI3K/AKT/mTOC1 signaling and enhanced translational activity. Together, these data highlight a potential mechanistic link between epigenetic regulation and translational control in the context of phenotypic plasticity. While the precise molecular mechanisms remain to be fully elucidated, our results raise the possibility that EZH2 contributes to plasticity not only through chromatin remodeling but also by influencing the protein synthesis machinery. These insights point to components of the translational apparatus as candidate therapeutic vulnerabilities in EZH2-driven malignancies.

Motivated by these findings, we sought to identify rational combination therapies targeting the translational machinery, particularly in the context of NEPC, where plasticity-driven resistance remains a clinical challenge. Our previous data had demonstrated the efficacy of combining the HDAC inhibitor Panobinostat and the dual PI3K/mTOR inhibitor BEZ235 in models of CRPC with PTEN deficiency (54). Similar results were observed using Fimepinostat, a dual HDAC/PI3K inhibitor in NEPC preclinical models (55). Moreover, the dual PI3K/mTOR inhibitor Gedatolisib has been shown to present greater anti-proliferative activity in PCa cells than single target inhibitors (56), and combination strategies that include PI3K/AKT/mTOR and AR-targeting therapies have shown superior efficacy in androgen-sensitive PCa cell lines and PDX models (57). Using independent mouse models representing NEPC generated through co-deletion of *Pten and Rb1*(3) or *Pten and Tp53* (58), along with a human NEPC model (37) we demonstrated that monotherapy with EZH2 or PI3K/mTOR inhibitor provided limited anti-tumor benefit in both intact and castrate settings. Modest responses to the combination therapy were seen in intact animals, but striking anti-tumor efficacy was observed under castrate conditions. Because tumor progression in these models involves a shift from AR-positive CRPC to AR-negative NEPC, we propose that EZH2 and PI3K/mTOR inhibitor co-treatment is particularly effective in castrate conditions due to EZH2 inhibition restoring AR expression and function. Additionally, we observed that EZH2 inhibition suppressed inflammatory signaling, consistent with prior work suggesting that inflammatory reprogramming is required for lineage transition from prostate adenocarcinoma to NEPC (5). Mechanistically, EZH2 has been shown to interact with RNA-binding proteins (RBPs) such as FBL (fibrillarin) or LIN28, influencing rRNA processing and the translation of specific oncogenic transcripts, including MYC, and MYCN (59). These interactions enhance protein synthesis by promoting ribosome biogenesis or promoting mRNA stability. We had recently shown that loss of the RNA-binding protein tristetraprolin (TTP), a key suppressor of inflammation, promotes tumor plasticity and resistance to enzalutamide in a mouse model of prostate adenocarcinoma (38). In this study, we observed that Crispr knockout of *Zfp36* (encoding TTP) impaired the ability of EZH2 inhibition to repress inflammatory signaling and to restore AR and PI3K/AKT/mTORC1 pathway activity. Notably, in the absence of TTP, combined inhibition of EZH2 and PI3K/mTOR signaling failed to elicit anti-tumor effects in both DKO and PPKO tumor cells, particularly under castrate conditions. These findings suggest that TTP may serve as a critical intermediary enabling the therapeutic response to EZH2 and PI3K/mTOR co-inhibition.

In several cancer types, such as small cell lung cancer, it has been reported that alterations in chromatin remodeling result in transcriptional dysregulation, driving tumor cells toward a highly plastic state that leads to metastatic tumors (60–62). In PCa, it has been previously demonstrated that there are unique DNA methylation landscapes defining the transition from adenocarcinoma to NEPC (2). Further, it is shown that defined histone methylation (H3K4me3 and H3K27me3) landscapes emerged when the LNCaP adenocarcinoma cell line was engineered to overexpress MYCN, a known NEPC-related transcription factor (63, 64). The association of retinoblastoma protein with chromatin is documented (65), and demonstrates promiscuous activity, where it binds at promoters and enhancers in a cell-type–specific manner. In the absence of RB1, we observe a significant remodeling of H3K4me3 and H3K27me3 histone marks associated with promoter regions and activation of key genes implicated in NEPC, including Notch1 (66), Tacstd2 (67), and MYCN (63), highlighting RB1’s critical role in maintaining chromatin architecture. These findings align with previous reports showing that EZH2 inhibition can induce a poised chromatin state (13), further supporting its role in epigenetic regulation of plasticity.

Collectively, our study demonstrates the control by EZH2 catalytic function in cellular plasticity and the numerous cell state transitions that converge during progression of adenocarcinoma to NEPC. Inhibition of EZH2 results in molecular changes which are therapeutically targetable and highlight their potential to treat patients whose tumors have evolved due to plasticity. In line with these findings, a previous paper in triple negative breast cancer, reported that the dual inhibition of AKT and EZH2 induces differentiation from an aggressive basal-like cells to a more differentiated, luminal-like phenotype (68). Our findings extend these insights to prostate cancer, underscoring the therapeutic promise of levering epigenetic reprogramming to uncover and exploit convergent vulnerabilities in translational regulation across heterogeneous and treatment-resistant cancer subtypes.

## Materials and Methods

### Study design

The study aimed to investigate the role of the epigenetic regulator EZH2 in driving phenotypic plasticity during the transition of adenocarcinoma to a NEPC state. Using multi-omics approaches, we determined that EZH2 regulates a multilineage cell state, which is dependent on the activation of translation and regulation of RNA maturation. The therapeutic potential of EZH2 inhibition was evaluated as monotherapy and in combination with a PI3K/mTORC1inhibitor. Expression of ZFP36 was examined using H. Beltran patient cohort (33 CRPC-Adeno, and 12 NEPC cases), which sufficient sample size to analyze statistical significance. Mice for in vivo studies were randomly assigned to control or treatment groups on the basis of tumor size, and there were no significant differences at the start of the study. Data collection was terminated when mice met the criteria in accordance with Institutional Animal Care and Use Committee (IACUC) guidelines or when predetermined treatment schedule was completed. All experiments were blinded where possible. For in vitro experiments, at least three independent experiments were performed.

### Animal ethics statement

All animal breeding and experiments were approved by and performed in accordance with the guidelines of the Cedars Sinai Medical Center, Institutional Animal Care and Use Committee (Animal protocol #9505).

### Cell culture, reagents, and drug response assays

Murine *Pten:Rb1* knock-out (DKO) 2D cell lines were generated from previously described genetically engineered mouse models (3)*. Pten:Tp53* knock-out (PPKO) 2D cell lines were kindly provided by Dr. Zongbing You (36). DKO and PPKO cell lines were cultured high-glucose DMEM with 1% P/S and either 10% FBS or CSS. Human WCM154 3D organoids have been described previously (2). For drug sensitivity assays, 500 or 350 cells/well for DKO and PPKO respectively were seeded in 3D in a 96 well plate pre-coated with collagen type I in suspension with 30% matrigel and treated with indicated concentrations of LY3023414 (LY-PK, class I PI3K/mTOR/DNA-PK inhibitor), and LY3346149 (LY-EZ, EZH2 inhibitor), and EPZ6438 (EPZ, EZH2 inhibitor) for 72 hours. Spheroid growth was assessed by CellTiter-Glo (Promega) assay according to manufactures instructions. Luminescence intensity of each well was normalized to the average of the control wells on the same plate to calculate relative cell viability values. For synergy analysis, relative cell viability measurements were analyzed using the web-based tool SynergyFinder (69). Bliss Synergy summary scores were derived from the average of the synergy scores across the entire dose–response landscape. Bliss analysis assume that the given drugs produce their anticancer effect by targeting different pathways, and that these pathways have no mechanistic connection other than the response outcome. Data visualization and statistical testing was performed using Prism 8 (GraphPad Software). Antibodies for ChIP-seq were H3K4me3 (C42D8) and H3K27me3 (C36B11) and purchased from Cell Signaling Technology. For immunohistochemistry, above histone antibodies were used, Ki67 (D3B5), and phospho-H2AX (20E3) antibodies were purchased from Cell Signaling Technology. For western blotting, androgen receptor antibody was purchased from Abcam and GAPDH from Cell Signaling Technology.

### Single-nuclear RNA sequencing (snRNA-seq)

Nuclei were isolated from fresh frozen mouse prostatic tumors using a method modified from a recent single-nuclei RNA-sequencing (snRNAseq) study. The ST-SB buffer from that study was modified by removing Tween-20 and supplementing with 0.04 U/μL Protector RNase Inhibitor (Roche). All sample manipulation was performed on wet ice with wide-bore pipet tips (Rainin) and all centrifugations were performed with a swinging bucket rotor maintained at 4 °C for 5 min at 850 × g. In brief, the frozen tissue was transferred onto a plate on dry ice and crushed into ≤1 mm3 pieces. This was then transferred to a 2 mL dounce homogenizer (Kimble, cat: 885300-0002) on wet ice containing 1 mL of Nuclei EZ lysis buffer (Sigma, cat: NUC101). The tissue was then dunked approximately ×20 with Pestle A followed by ×20 with Pestle B. The lysis was then quenched by adding 1 mL of ST-SB. The sample was filtered through a pre-wetted 30 μm filter (Miltenyi Biotec, cat: 130-041-407) into a 15 mL conical tube. The homogenizer was rinsed 3× with 1 mL of ST-SB and this was transferred through the same 30 μm filter into the 15 mL conical tube. The sample was then centrifuged, the resulting supernatant removed, and the pellet resuspended with 500 μL of ST-SB. After, the sample was passed through a pre-wetted 20μm filter (Miltenyi Biotec, cat: 130-101-812) into a 1.5 mL protein lo-bind microcentrifuge tube (Eppendorf, cat: 022431081) and centrifuged. Nuclei concentration was quantified by mixing an aliquot of the sample with DAPI at a final concentration of 0.025 mg/mL in H2O. Samples were finally processed according to 10×Genomics protocol for the 3’ v3.1 assay and were super-loaded to target of 20,000 nuclei recovery. Single-cell RNA-Seq libraries were prepared per the 10x Genomics Single Cell 3′ v3.1 Reagent Kits User Guide using the Chromium Controller, X, or Connect (10x Genomics, Pleasanton, California). Barcoded sequencing libraries were quantified by quantitative PCR using the Collibri Library Quantification Kit (Thermo Fisher Scientific, Waltham, MA). Fragment analysis was performed on the 4200 TapeStation (Agilent Technologies, Santa Clara, California). Libraries were sequenced on a NovaSeq 6000 (Illumina, San Diego, CA) as per the Single Cell 3′ v3.1 Reagent Kits User Guide, with a sequencing depth of ∼40,000 reads/cell. The demultiplexed raw reads were aligned to the transcriptome using STAR (version 2.5.1) (Dobin et al., 2013) with default parameters, using mouse mm10 transcriptome reference from Ensembl version 84 annotation, containing all protein coding and long non-coding RNA genes. Expression counts for each gene in all samples were collapsed and normalized to unique molecular identifier (UMI) counts using Cell Ranger software version 4.0.0 (10X Genomics). The result is a large digital expression matrix with cell barcodes as rows and gene identities as columns.

### Chromatin immunoprecipitation with sequencing (ChIP-seq)

Fresh-frozen prostatic tumor tissue was pulverized (Cryoprep Pulvrizer, Covaris), resuspended in PBS + 1% formaldehyde, and incubated at room temperature for 20 minutes. Fixation was stopped by addition of 0.125 M glycine (final concentration) for 15 minutes at room temperature, then washing in ice cold PBS + EDTA-free protease inhibitor cocktail (PIC; #04693132001, Roche). Chromatin was isolated from biological triplicates by the addition of lysis buffer (0.1% SDS, 1% Triton X-100, 10 mM Tris-HCl (pH 7.4), 1 mM EDTA (pH 8.0), 0.1% NaDOC, 0.13 M NaCl, 1X PIC) + sonication buffer (0.25% sarkosyl, 1 mM DTT) to the samples, which were maintained on ice for 30 minutes. Lysates were sonicated (E210 Focused-ultrasonicator, Covaris) and the DNA was sheared to an average length of ∼200-500 bp. Genomic DNA (input) was isolated by treating sheared chromatin samples with RNase (30 minutes at 37°C), proteinase K (30 minutes at 55°C) and de-crosslinking buffer (1% SDS, 100 mM NaHCO3 (final concentration), 6-16 hours at 65°C), followed by purification (#28008, Qiagen). DNA was quantified on a NanoDrop spectrophotometer, using the Quant-iT High-Sensitivity dsDNA Assay Kit (#Q33120, Thermo Fisher Scientific). On ice, H3K27me3 (5 µl, #07-449, Millipore) or H3K4me3 (5 µl, #17-614, Millipore) antibodies were conjugated to a mix of washed Dynalbeads protein A and G (Thermo Fisher Scientific) and incubated on a rotator (overnight at 4°C) with 1.3 µg of chromatin. ChIP’ed complexes were washed, sequentially treated with RNase (30 minutes at 37°C), proteinase K (30 minutes at 55°C), de-crosslinking buffer (1% SDS, 100 mM NaHCO3 (final concentration), 6-16 hours at 65°C), and purified (#28008, Qiagen). The concentration and size distribution of the immunoprecipitated DNA was measured using the Bioanalyzer High Sensitivity DNA kit (#5067-4626, Agilent). Dana-Farber Cancer Institute Molecular Biology Core Facilities prepared libraries from 2 ng of DNA, using the ThruPLEX DNA-seq kit (#R400427, Rubicon Genomics), according to the manufacturer’s protocol; finished libraries were quantified by the Qubit dsDNA High-Sensitivity Assay Kit (#32854, Thermo Fisher Scientific), by an Agilent TapeStation 2200 system using D1000 ScreenTape (# 5067-5582, Agilent), and by RT-qPCR using the KAPA library quantification kit (# KK4835, Kapa Biosystems), according to the manufacturers’ protocols; ChIP-seq libraries were uniquely indexed in equimolar ratios, and sequenced to a target depth of 40M reads on an Illumina NextSeq500 run, with single-end 75bp reads. BWA (version 0.6.1) was used to align the ChIP-seq datasets to build version NCB38/MM10 of the mouse genome. Alignments were performed using default parameters that preserved reads mapping uniquely to the genome without mismatches. Bam files were concatenated to sum the biological replicates of each state and bigwiggle files were calculated for comparing the different states.

### Crispr Screen

To carry out the pooled genome-wide Crispr screen, 1 × 10^8^ DKO-Cas9 cells were infected with the pooled lentiviral Brie library (lenti-Guide-Puro backbone) system (28) at a multiplicity of infection of 0.5. Following 3 days of puromycin selection, half the cells were stored as day 0 control samples, and the remaining cells were cultured in DMSO or EPZ6438 (1μM) for an additional 2 weeks. PCR was carried using genomic DNA to construct the sequencing library which were then sequenced at approximately 30-35 million reads to achieve around 300x average coverage over the Crispr library. Data analysis was conducted using MAGeCK (30).

### RIME Assay

#### Chromatin Immunoprecipitation

Chromatin was isolated from fixed DKO and SKO cells by the addition of lysis buffer, followed by disruption with a Dounce homogenizer. Lysates were sonicated and the DNA sheared to an average length of 300-500 bp. Genomic DNA (Input) was prepared by treating aliquots of chromatin with RNase, proteinase K and heat for de-crosslinking, followed by ethanol precipitation. Pellets were resuspended and the resulting DNA was quantified on a NanoDrop spectrophotometer. Extrapolation to the original chromatin volume allowed quantitation of the total chromatin yield. An aliquot of chromatin (150 ug) was precleared with protein G agarose beads (Invitrogen). Proteins of interest were immunoprecipitated using 15ul of antibody against EZH2 (Active Motif, Cat# 39901) and protein G magnetic beads. Protein complexes were washed then trypsin was used to remove the immunoprecipitate from beads and digested the protein sample. Protein digests were separated from the beads and purified using a C18 spin column (Harvard Apparatus). The peptides were vacuum dried using a speedvac.

#### Mass Spectrometry

Digested peptides were analyzed by LC-MS/MS on a Thermo Scientific Q Exactive Orbitrap Mass spectrometer linked to Dionex Ultimate 3000 HPLC (Thermo Scientific) and a nanospray FlexTM ion source. The digested peptides were loaded directly onto the separation column Waters BEH C18, 75-micron × 100 mm, 130Å 1.7u particle size. Peptides were eluted using a 100-minute gradient with a flow rate of 323 nl/min. An MS survey scan was obtained for the m/z range 340-1600, MS/MS spectra were acquired using a top 15 method, where the top 15 ions in the MS spectra were subjected to HCD (High Energy Collisional Dissociation). An isolation mass window of 1.6 m/z was for the precursor ion selection, and normalized collision energy of 27% was used for fragmentation. A 20 second duration was used for the dynamic exclusion.

#### Database Searching

Tandem mass spectra were extracted and analyzed by PEAKS Studio version 8 built 20. Charge state deconvolution and deisotoping were not performed. Database consisted of the Uniprot database (version 180508, 71,771 curated entries) and the cRAP database of common laboratory contaminants (www.thegpm.org/crap; 114 entries). Database was searched with a fragment ion mass tolerance of 0.02 Da and a parent ion tolerance of 10 PPM. Post-translational variable modifications consisted of methionine oxidation, asparagine and glutamine deamidation.

#### Criteria for Protein Identification

Peaks studio built-in decoy sequencing and FDR determination was used to validate MS/MS based peptide and the parsimony rules for protein identifications. A threshold of the −10*logp (p-value) of 20 or greater was applied for the peptide identifications. The weighted sum of 9 parameters for peptide scoring are converted to a p-value which represent the probability of a false identification. Protein identifications were accepted if they could pass the - 10logp of 20 and contained at least 1 identified unique peptide. Proteins that contained similar peptides and could not be differentiated based on MS/MS analysis alone were grouped to satisfy the principles of parsimony. Proteins sharing significant peptide evidence were grouped into protein groups.

#### List Filtering

Protein coverage percentage is observed in the raw data file. Final list generation was done by taking all proteins with a spectral count of five and above from each replicate reaction and comparing them in a venn-diagram against IgG control replicates. Proteins unique to both experimental replicates were then applied to the PANTHER database for protein ontology results. Database *Searching*: Tandem mass spectra were extracted and analyzed by PEAKS Studio version 8 built 20. Charge state deconvolution and deisotoping were not performed. Database consisted of the Uniprot database (version 180508, 71,771 curated entries) and the cRAP database of common laboratory contaminants (www.thegpm.org/crap; 114 entries). Database was searched with a fragment ion mass tolerance of 0.02 Da and a parent ion tolerance of 10 PPM. Post-translational variable modifications consisted of methionine oxidation, asparagine and glutamine deamidation.

### In vivo therapy experiments

#### Mouse transplant models

Experiments were carried out using 8-week-old male C57BL/6N mice (Charles River Labs). Mice were subcutaneously injected with DKO or PPKO cells contained in matrigel/PBS (50/50). At time of tumor implantation, mice were either surgically castrated (castrated) or sham castrated (intact). Therapy started when tumors measured approximately 20mm^2^. Therapy consisted of daily of LY-PK (15mg/kg) and/or LY-EZ (20mg/kg) by intraperitoneal (IP) injection.

#### Human transplant model

Experiments were carried out using 8-week-old male NOD.CB17-Prkdcscid/NCrCrl mice (Charles River Labs). Mice were subcutaneously injected with WCM154 cells contained in matrigel/PBS (50/50). Mice were castrated 7 days before tumor transplant. Therapy started when tumors measured approximately 20mm^2^. Therapy consisted of daily of LY-PK (15mg/kg) and/or LY-EZ (20mg/kg) by intraperitoneal (IP) injection, with or without enzalutamide (30mg/kg) by oral gavage.

For all in vivo experiments, tumors were measured three times weekly by caliper measurements. Treatment toxicities were assessed by body weight, decreased food consumption, signs of dehydration, hunching, ruffled fur appearance, inactivity, or nonresponsive behavior.

#### In vivo immunohistochemistry

Four-(4)-μm-thick, formalin-fixed, paraffin-embedded sections were prepared using standard sodium citrate antigen-retrieval methods. Primary antibodies were diluted in 1.25% horse serum/ PBS and incubated overnight in a humidified chamber at 4 °C. IHC staining was carried out using the ImmPRESS horseradish peroxidase anti-mouse IgG (peroxidase) Polymer Detection Kit (Vector Laboratories) following the manufacturer’s instructions for visualization. Sections were incubated with primary antibody in a humidified chamber at 4 °C overnight, washed in PBS-T and cover slipped with VECTASHIELD Antifade Mounting Medium with DAPI (Vector Laboratories).

### Zfp36 CRISPR-knockout cell lines and RNA-seq

For CRISPR/Cas9-mediated knockout cell line generation in DKO and PPKO cells, guide RNA (gRNA) sequences CATGACCTGTCATCCGACCA and AAGCGGGCGTTGTCGCTACG targeting murine *Zfp36*, were cloned into the lenti-CRISPR/Cas9v2 vector (Addgene, #52961) according to the Zhang lab protocol. The scrambled gRNA sequence CACCGCGTGATGGTCTCGATTGAGT was used as a negative control. Viral infection was performed as described by the RNAi consortium (Broad Institute) laboratory protocol “Lentivirus production of shRNA or ORF-pLX clones,” and single clones were isolated following puromycin selection (2 μg/mL; Sigma-Aldrich #P8833). The DKO control (DKO Ctl) and DKO:Zfp36 KO (DKO KO) cells were treated with the indicated doses of LY3023414 (LY-PK, dual PI3K/mTORC1/2 inhibitor), LY3346149 (LY-EZ, EZH2 inhibitor), and EPZ6438 (EPZ, EZH2 inhibitor) for 96 hours. RNA-seq data were aligned with STAR to mouse reference genome mm10 (GRCm38), quantified using RSEM and normalized. Gene set enrichment analysis (GSEA) was performed in GenePattern (genepattern.org). Genotypes were compared as described, using 10,000 gene set permutations to generate normalized enrichments scores, with FDR q-value <0.25 considered significant.

### Western Blot

Sub-confluent treated cells were washed twice with cold PBS, trypsinized and then lysed in Pierce RIPA buffer (Thermo Fisher Scientific, 89900) with PhosSTOP™ inhibitor cocktail (Sigma Aldrich, PHOSS-RO) at 4°C for 30 mins. Protein concentrations were measured by a Pierce BCA Protein Assay Kit (Thermo Fisher Scientific, 23225). Proteins were separated by SDS-PAGE (10% Mini-PROTEAN® TGX™ Precast Gel, Biorad, 4561036) and transferred to a PVDF membrane (Bio-Rad, 1620177). The membrane was blocked 3% BSA in TBST for 1 hour at room temperature and then blotted with primary antibodies overnight at 4°C. Blots were washed 3 x 5 minutes with TBST. The blots were then incubated with fluorescent-conjugated secondary antibody (Bio-Rad) 1 hour at room temperature and washed 3 x 5 minutes with TBST. Proteins were visualized with a ChemiDoc MP fluorescent imager (Bio-Rad). Bio-Rad Image Lab analysis software was used to quantify protein band density. All western blots were repeated twice.

### Intracellular antibody staining for flow cytometry

4-6 million cells were fixed 15 min with 4% PFA and permeabilized with 0.1% Triton by 30 minutes. After two washes with permeabilization buffer, cells were resuspended in FACS buffer containing antibody against Zfp36 (LS-B5606, LSBio), TSC2 (D93F12, Cell Signaling), p-S6K (9205, Cell Signaling), and p4E-BP1 (2855, Cell Signaling) and incubated 30 minutes at room temperature. After two washes cells were resuspended in FACS buffer containing the secondary antibody (Biolegend 406410). After two additional washes with permeabilization buffer, cells were resuspended in FACS buffer (1% of FCS+2mM EDTA in PBS).

## Data, Materials, and Software Availability

All data needed to evaluate the conclusions are present in the paper and/or the Supplementary Materials. Code for snATACseq analysis is available at Github repository on demand.

## Acknowledgments

We thank the Center for Functional Cancer Epigenetics at Dana-Farber Cancer Institute, and the Center for Bioinformatics and Functional Genomics at Cedars Sinai Medical Center for sequencing and data analysis assistance. This study was supported from a Department of Defense Early Career Investigator Award (W81XWH-22-PRCP-EIRA to BG and W81XWH1910305 to KLM), and the NCI (R01CA252468 to LE). The contents of this publication are the sole responsibility of the author(s) and do not necessarily reflect the views, opinions or policies of Uniformed Services University of the Health Sciences (USUHS), The Henry M. Jackson Foundation for the Advancement of Military Medicine, Inc, the Department of Defense (DoD) or the Departments of the Army, Navy, or Air Force. Mention of trade names, commercial products, or organizations does not imply endorsement by the U.S. Government.

The contents of this publication are the sole responsibility of the author(s) and do not necessarily reflect the views, opinions or policies of Uniformed Services University of the Health Sciences (USUHS), The Henry M. Jackson Foundation for the Advancement of Military Medicine, Inc, the Department of Defense (DoD) or the Departments of the Army, Navy, or Air Force. Mention of trade names, commercial products, or organizations does not imply endorsement by the U.S. Government

## Author Contributions

Conceptualization: LE, BG, KLM

Methodology: LE, BG, KLM, TN, NB, FD, XQ, HWL, MB, SRVK, JTP, DPL

Investigation: LE, BG, KLM, DLB, TN, NB, FD, SRVK, JTP, DPL

Analysis: LE, BG, KLM, DLB, TN, NB, AAS, SD, FD, SRVK, JTP, DPL

Visualization: LE, BG, KLM, DLB, TN, NB, FD, MLF, SRVK, JTP, DPL

Funding acquisition: LE, BG, KLM

Project administration: LE, BG, KLM

Supervision: LE

Writing – original draft: LE, BG, KLM, DLB, TN, NB, FD, MLF, SRVK, JTP, DPL

Writing – review & editing: LE, BG, KLM, DLB, TN, NB, FD, MLF, SRVK, JTP, DPL

Writing/Editing: LE, BG, KLM, DLB, TN, NB, FD, MLF, SRVK, JTP, DPL

---

**Supplementary figure 1:**
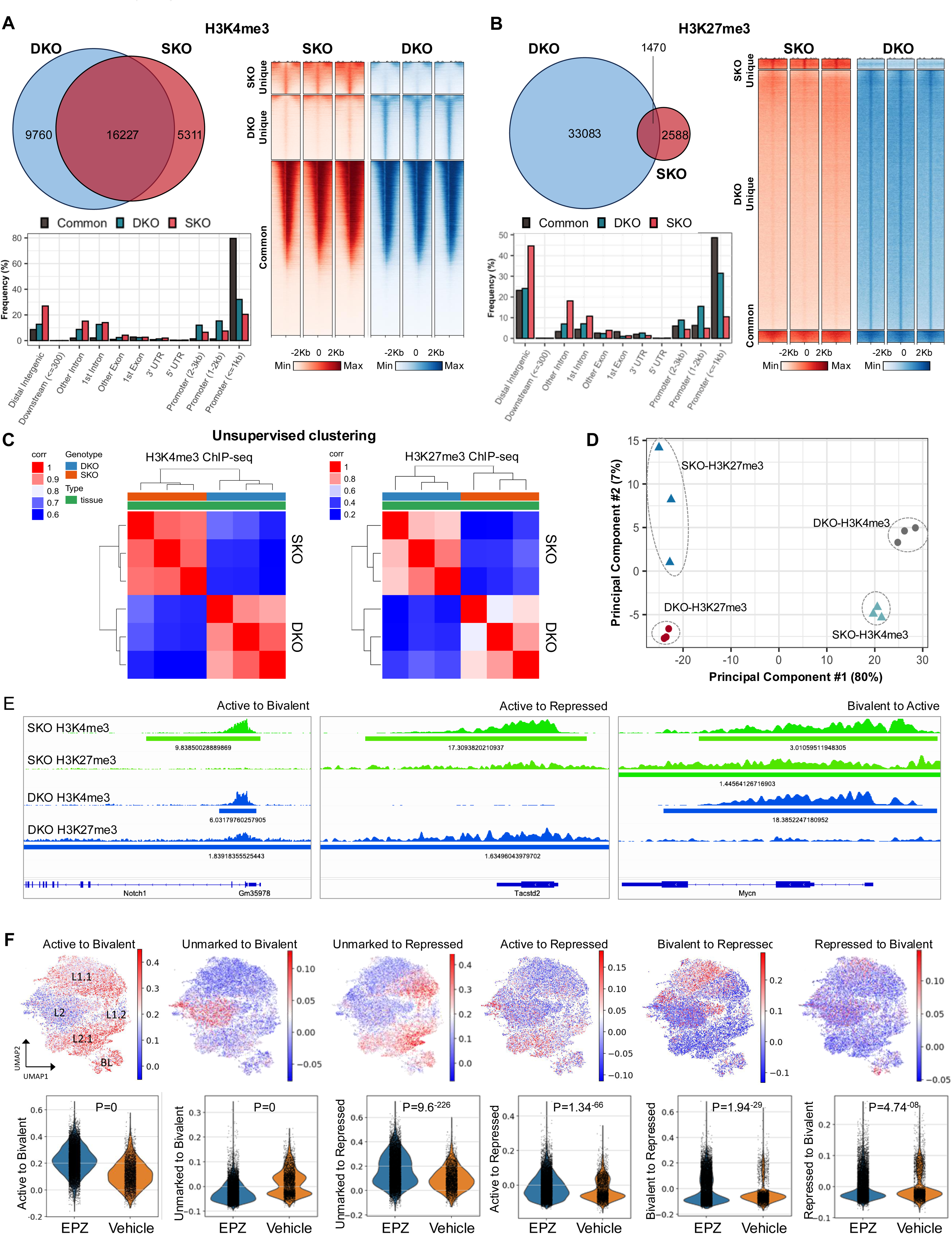
EZH2 coordinates multi-lineage transitional cell states mediated by RB1 loss. **A**. Venn diagram (left) and tornado plot (right) illustrating the 31,298 ChIP-seq peaks for H3K4me3 in *PbCre:Pten KO* (SKO) and *PbCre:Pten:Rb1 KO* (DKO) tumors. Bar plot (bottom left) indicates H3K4me3 peak genome-wide peak distribution. 52% of the peaks were shared between SKO and DKO, 17% and 31% were considered unique in SKO and DKO respectively. The analysis of exact genomic location of H3K4me3 is illustrated in the bars plot and showed that most of the genomic locations are at promoter regions. **B.** Venn diagram (left) and tornado plot (right) illustrating the 37,141 ChIP-seq peaks for H3K27me3 in SKO and DKO tumors. 4% of the peaks were shared between SKO and DKO, 7% and 89% were considered unique in SKO and DKO respectively. The analysis of exact genomic location of H3K27me3 is illustrated in the bars plot and showed that most of the genomic locations are at promoter regions. **C.** Hierarchical clustering of SKO and DKO tumors based on sample-to-sample correlation of H3K4me3 and H3K27me3 peaks representing promoter occupancy. **D.** Two-dimensional principal component analysis (PCA) of H3K4me3 and H3K27me3 peaks representing promoter occupancy **E.** Gene track examples chromatin remodeling in Notch1 and Tacstd2 which move from an active to bivalent or repressed state respectively and Mycn from a bivalent to active state. **F.** UMAP representing the genes transitioning from Active to Bivalent, Unmarked to Bivalent, unmarked to repressed, active to repressed, bivalent to repressed and repressed to bivalent from SKO to DKO tumors based on the H3K4me3 and H3K27me3 ChIP-seq analysis (top). Violin plots of significantly enriched gene sets in DKO upon vehicle or EPZ treatment. FDR < 0.01, abs (NES) > 1 where NES is the normalized enrichment score.

**Supplementary Figure 2:**
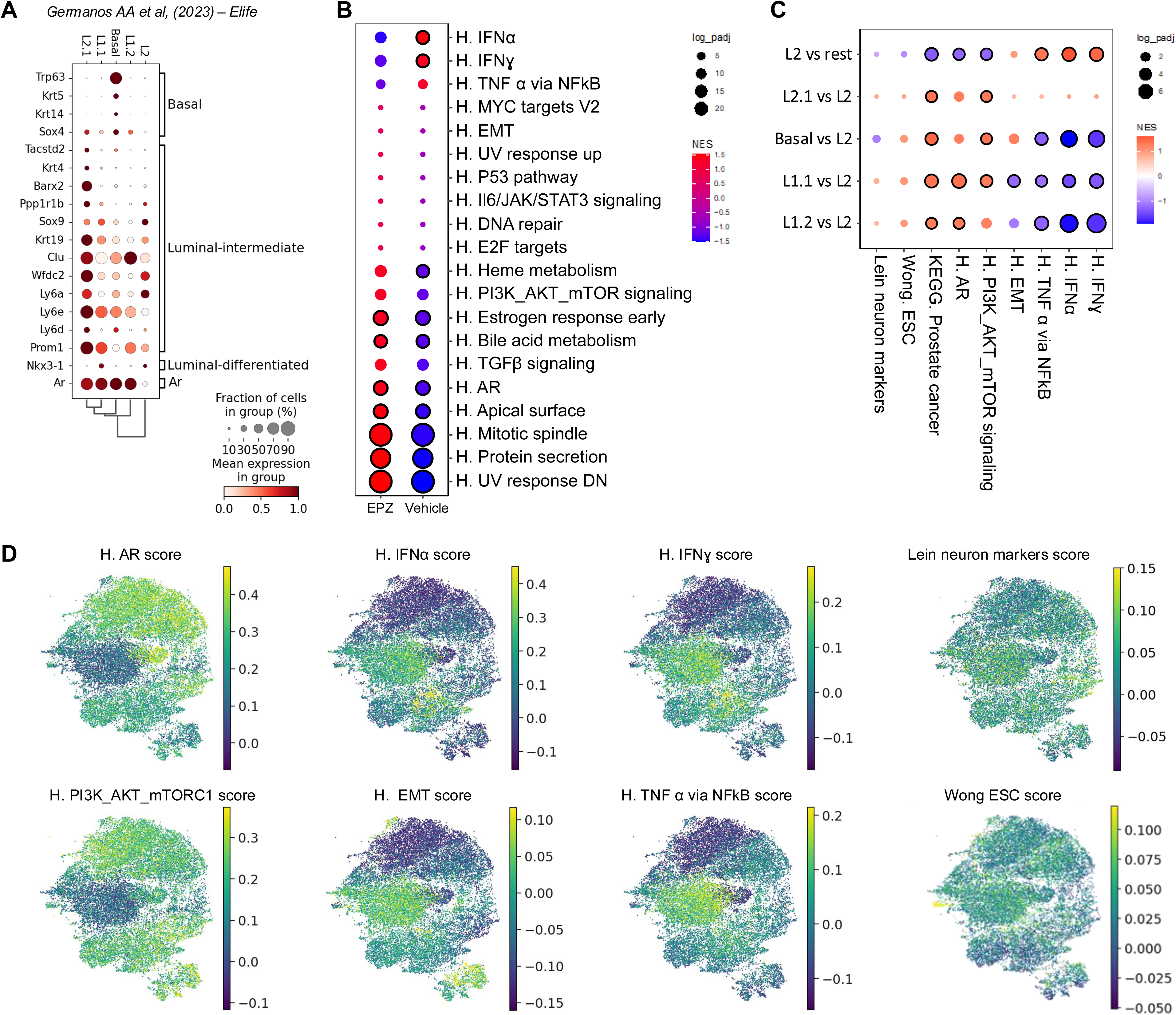
**A**. Dot plot of canonical marker genes for each cell type. Hue of red indicates average normalized gene expression level and size of dot indicates percentage of nonzero gene expression for each cell population. **B.** Dot plot of the Gene Set Enrichment Analysis (GSEA) of transcripts upregulated in EPZ treated mice compared with vehicle. **C.** Dot plot of the GSEA of transcripts upregulated in each epithelial sub-cluster compared with L2. **D.** UMAP of the androgen signaling (NELSON_ANDROGEN_ SIGNALING_UP), epithelial mesenchymal transition (EMT), interferon alpha response (IFNα), interferon gamma response (IFNɣ), PI3K-AKT-MTOR signaling and TNFα signaling via NFkB, embryonic stem cells.

**Supplementary Figure 3:**
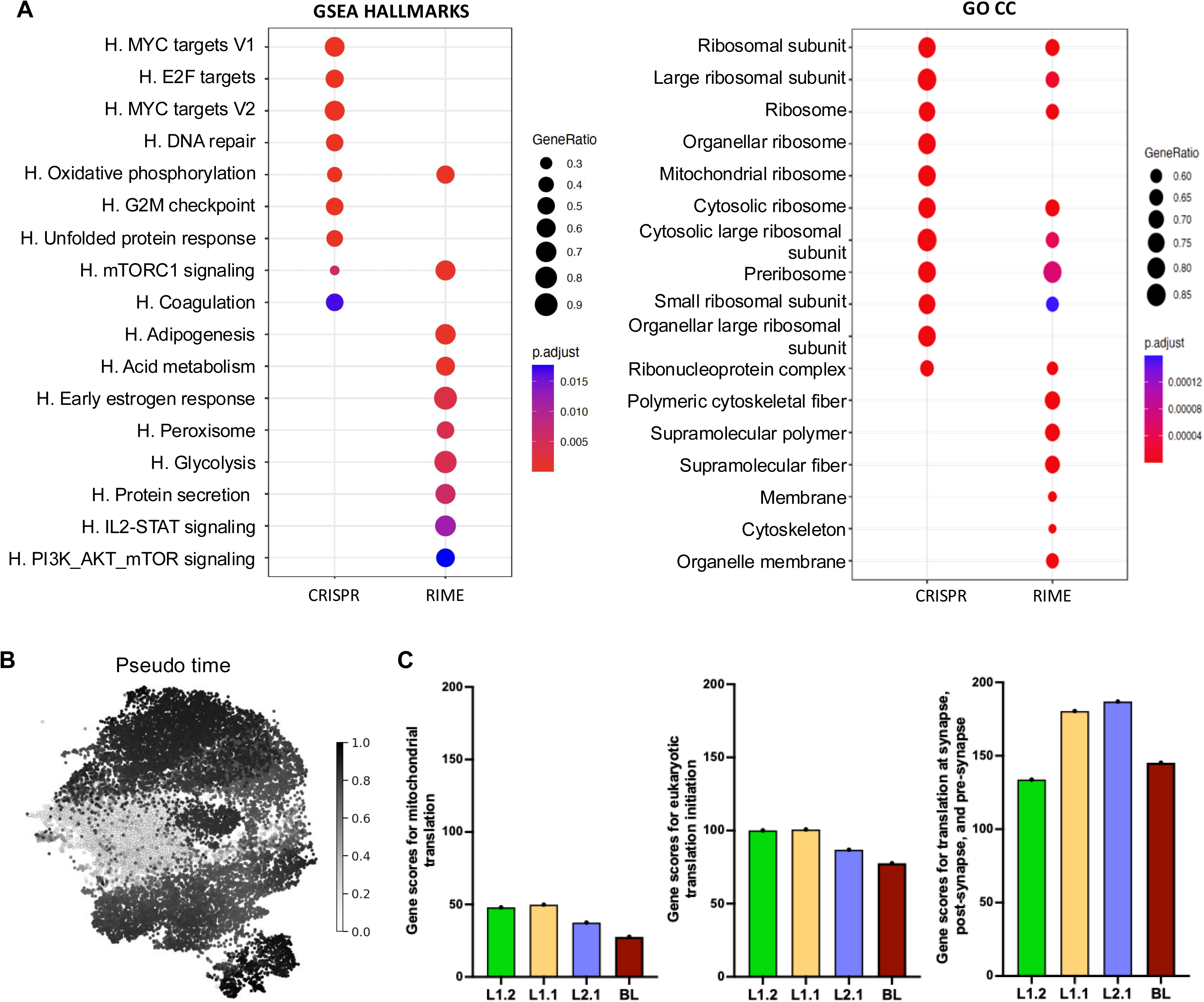
**A.** Dot plot representing the Gene Set Enrichment Analysis (GSEA) Hallmarks pathway and gene ontology cellular component analysis demonstrate overlay of common pathways and process’s from Crispr screening and RIME analysis for DKO and SKO cells. The blue and red color represents the p-values across all cells, and the size of the dot corresponds to the percentage of cells expressing the characteristic for each pathway. **B.** Pseudo time analysis of epithelial cells in vehicle and EPZ treated mice. **C.** Bar plots showing the signature related with translation mechanisms in L2 sub-cluster versus other sub-clusters (L2.1, L1.1, L1.2 and BL). The analyzed signatures include genes related with processing of capped intron pre-mRNA, eukaryotic translation initiation, and mitochondrial translation.

**Supplementary figure 4:**
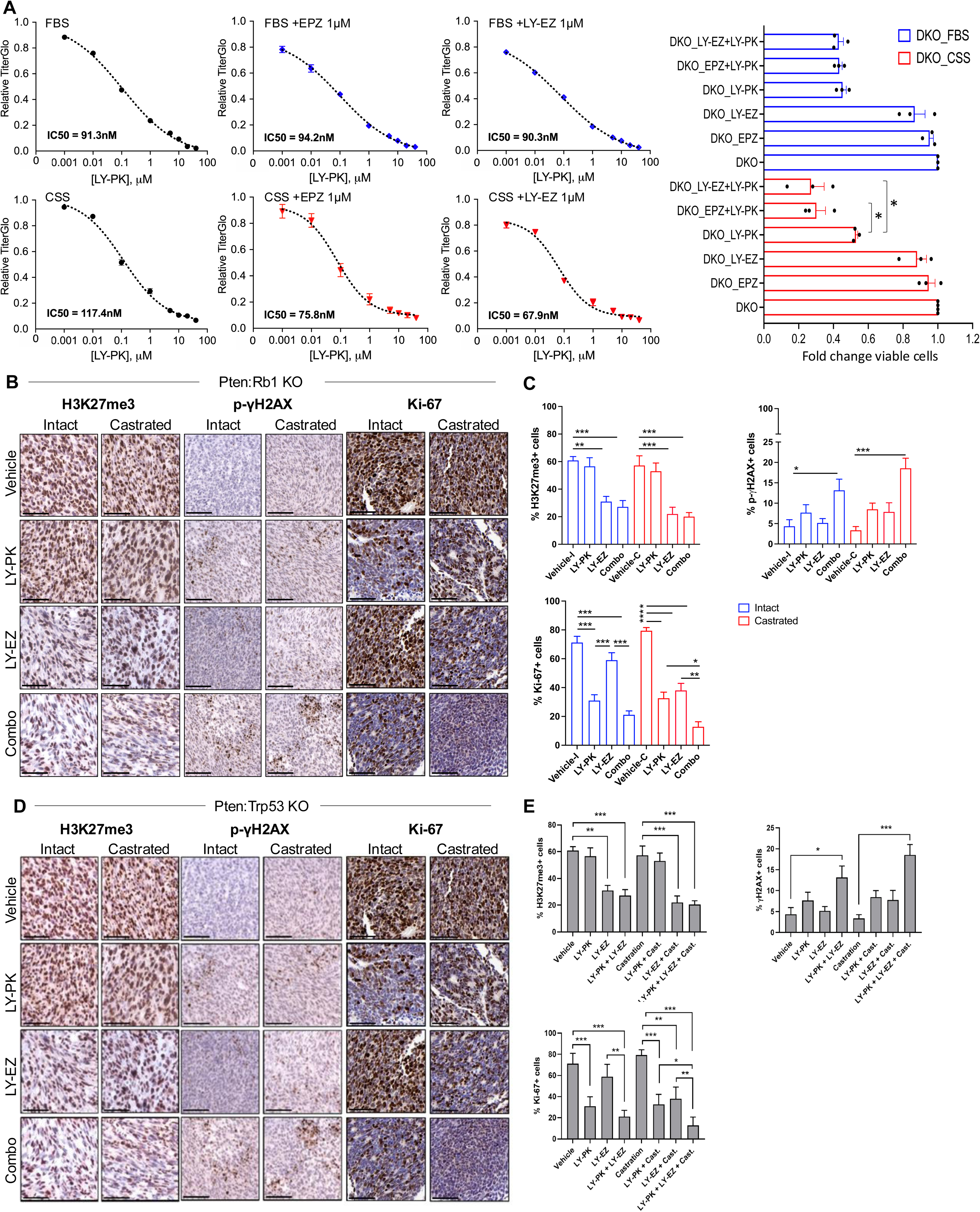
**A.** Titre glo and cell viability fold change from DKO spheroids cultures in FBS and CSS with a LY-PK dose curve and 1µM of EPZ or LY-EZ. **B-E.** H3K27me3, p-γH2AX and Ki-67 IHC staining in murine DKO and PPKO tumors from the in vivo study, and the corresponding quantification of the percentage of positive cells (n = 5 mice per treatment group, +/-1SD).

**Supplementary figure 5:**
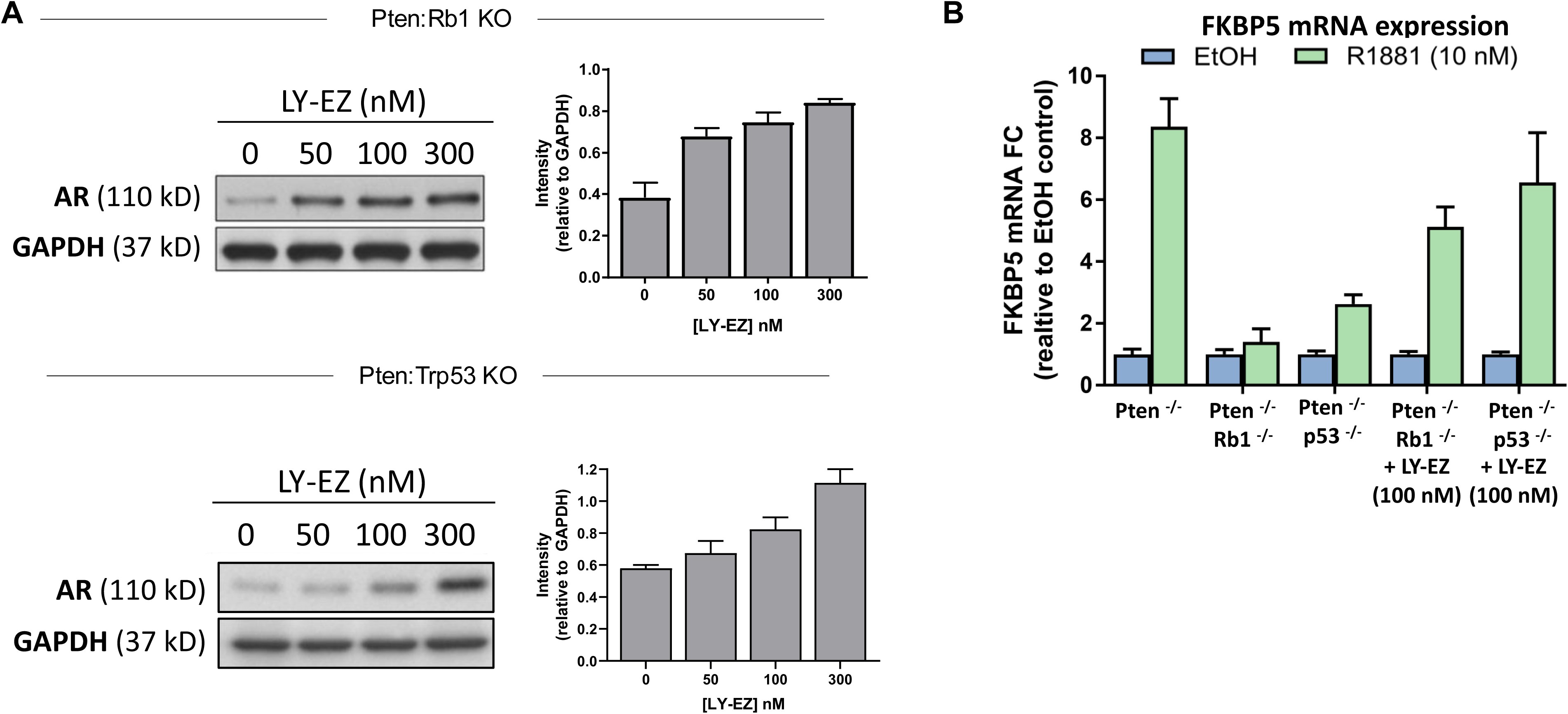
**A.** Western blot indicating AR protein levels and quantification in the DKO and PPKO cells treated with DMSO or LY-EZ at different concentrations (indicated in the figure). **B.** Fold change of Fkbp5 in GEMM-derived DKO and PPKO cells treated with EtOH or R1881 (10 nM) stimulation for 24hrs, in triplicates.

**Supplementary figure 6:**
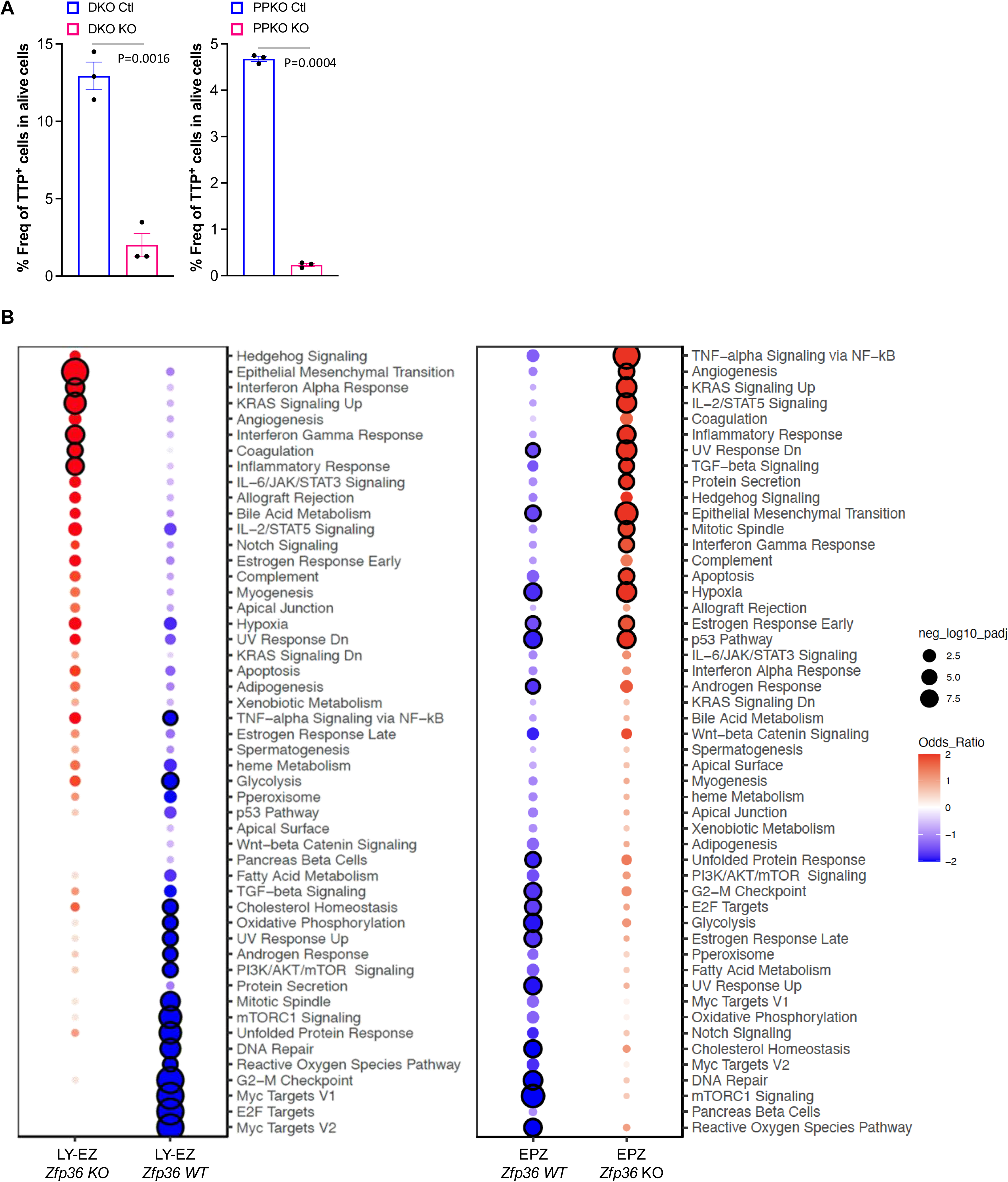
**A.** Bar plot representing the TTP flow cytometry analysis in DKO Ctl, PPKO Ctl, DKO *Zfp36* KO, PPKO *Zfp36* KO. **B**. Extended bubble plot demonstrating GSEA analysis of indicated gene signatures from RNAseq derived from DKO and cells treated with DMSO controls or the EZH2i (EPZ6438 or LY-EZ, 5μM) for 96 hours. Hue of red indicates average normalized gene signature level and size of dot indicates percentage of nonzero gene signature.

